# Sexual dimorphism and sex ratio bias in the dioecious willow *Salix purpurea* L

**DOI:** 10.1101/2020.04.05.026427

**Authors:** Fred E. Gouker, Craig H. Carlson, Junzhu Zou, Luke Evans, Chase R. Crowell, Christine D. Smart, Stephen P. DiFazio, Lawrence B. Smart

## Abstract

**Premise:** Sexual dimorphism in dioecious plant species is often not obvious or is absent. Dioecious species populations also often exhibit deviations from expected sex ratios. Previous studies on members of the Salicaceae family have shown strong, partial, and no sexual dimorphism. Some studies have shown sex-biased ratios in several *Salix* spp., however, *S. purpurea* has never been examined for evidence of sexual dimorphism or for the presence of sex-ratio bias, and therefore a comprehensive phenotypic study is needed to fill this knowledge gap.

**Methods:** This study examined a suite of morphological, phenological, physiological and wood composition traits from multi-environment and multi-year replicated field trials in a diversity panel of unrelated *S. purpurea* accessions and in full-sib F_1_ and F_2_ families produced through controlled cross pollinations to test for sexual dimorphism and sex ratio bias.

**Key Results:** Significant evidence of sexual dimorphism was found in vegetative traits with greater means for many traits in male genotypes compared to females across three populations of *S. purpurea*, measured across multiple years that were highly predictive of biomass yield. Male plants exhibited greater nitrogen accumulation under fertilizer amendment as measured by SPAD in the diversity panel, and males showed greater susceptibility to fungal infection by *Melampsora* spp in the F_2_ family. There were also consistent female-biased sex ratios in both the F_1_ and F_2_ families.

**Conclusions:** These results provide the first evidence of sexual dimorphism in *S. purpurea* and also confirm the prevalence of female-biased sex ratios previously found in other *Salix* species.

## INTRODUCTION

Dioecy is found in 4-10% of all flowering plants and is therefore much less common than the co-sexual states of monoecy and hermaphroditism (reviewed by Renner, 2014; Charlesworth, 2015; Sanderson *et al*., 2019), even if one includes the instances of subdioecy, which has been reported in 32 species in 21 families (Ehlers and Bataillon, 2007). Dioecy in flowering plants is therefore thought to have evolved from an ancestral co-sexual state since the first flowering plants evolved, approximately 124.6 million years ago (MYA) (Sun *et al*., 2002). Dioecy has evolved independently in many different plant families (reviewed by Renner, 2014) and even with different species in the same genus (Westergaard, 1958; Feng *et al*., 2020). In some plant lineages, homomorphic or hetromorphic sex chromosomes have evolved (Ming *et al*., 2007; Chen *et al*., 2016, reviewed by Charlesworth, 2015). Recent studies have identified sex-determination regions (SDRs) in dioecious plants including grape (*Vitis*), papaya (*Carica*), and persimmon (*Diospyros*), (Fechter *et al*., 2012; Wang *et al*., 2012; Akagi *et al*., 2014), in which the males are heterogametic, as well as other systems such as shrub willow (*Salix*) (Zhou *et al*., 2018), in which females are heterogametic.

Species of the Salicaceae family are almost all dioecious, including the woody perennial species in the genera *Populus* and *Salix*, which have clear morphological differences between staminate and pistillate catkins (Dickmann and Kuzovkina, 2008). Dioecy in the Salicaceae is thought to have evolved before the divergence of *Salix* and *Populus* approximately 65 MYA (Collinson, 1992; Tuskan *et al*., 2006). Multiple cytological studies of *Populus* suggest that Salicaceae have homomorphic chromosomes (Peto, 1938; van Buijtenen and Einspahr, 1959), and genetic mapping studies indicate that both male and female heterogametic systems are present in *Populus* (Pakull *et al*., 2009; Pakull *et al*., 2011; Tuskan *et al*., 2012; Kersten *et al*., 2014, Geraldes *et al*., 2015). The sex determining locus has been mapped to two different positions on Chr19 in different *Populus* species (Kersten *et al*., 2014; Geraldes *et al*., 2015). In willow species examined thus far, *S. viminalis, S. suchowensis*, and *S. purpurea*, females are the heterogametic sex (Semerikov *et al*., 2003; Hou *et al*., 2015; Pucholt *et al*., 2015; Chen *et al*., 2016; Zhou *et al*., 2018), in which the sex determination locus maps to Chr15 (Hou *et al*., 2015; Zhou *et al*., 2018) suggesting that the sex determination locus has translocated during recent evolution of *Populus* and *Salix*.

The relationship between sex chromosome evolution and sexual dimorphism and sex ratios are not yet fully understood. Primary sexual dimorphism refers to differences in the gametes produced, and in plants, this is expected to be controlled by sex-determining genes on a single sex chromosome, at least a portion of which is non-recombining (Charlesworth and Charlesworth, 1978). Secondary dimorphism includes other differences between males and females, such as morphology, physiology, and phenology (Charlesworth, 1999; Dawson and Geber, 1999; Delph, 1999), but in general sexual dimorphism in plants is less pronounced than in many animal systems, and evidence for dimorphism in secondary characteristics is scarce. Such differences may evolve when ecological and/or sexual selection lead to different fitness optima between sexes, and/or when physiological trade-offs occur due to the unequal energy costs of producing seeds versus pollen (Lewis, 1942; Arnold, 1994; Delph, 1999; Obeso, 2002). Females of woody dioecious plants typically produce less biomass than males, due to slower vegetative growth as a result of greater allocation to reproduction (Lewis, 1942; Lloyd and Webb, 1977; Obeso, 2002). Traits such as primary growth, production of secondary metabolites, and water use efficiency may be influenced by carbon resource allocation related to sex. Contrasting results have been found in *Salix*. In *S. sachalinensis* (syn *S. udensis*), no differences were detected in growth or mortality rates between males and females measured in a natural population over a three-year period (Ueno *et al*., 2007). Conversely, it was reported that drought tolerance and gas exchange rates differed between sexes in *S. glauca*, indicating dimorphism in physiological responses to abiotic stress with lower stomatal conductance (*g_s_*) and transpiration rates in males than in females when exposed to the same drought conditions (Dudley and Galen, 2007). However, the possibility of dimorphism has been studied for only a limited number of characteristics (Boecklen *et al*., 1994; Boecklen *et al*., 1990).

Another aspect of dimorphism in dioecious plants is sex ratio bias. Classical theories for sex ratios predict a 1:1 ratio if the expense per progeny is the same for both sexes (Fisher, 1930; Edwards, 2000). Theoretical models (reviewed by Delph, 1999; Obeso, 2002) suggest that sex ratio bias in natural populations could be due to differences in pollen and seed dispersal leading to different trade-offs between benefits and costs (Lloyd, 1982), other ecological factors (Barret *et al*., 2010), or to pollen competition in species with sex chromosomes (certation), such that X- or Y-bearing pollen is slow growing, or genetic transmission bias due to distorters of the sex chromosome segregation in meiosis (Taylor, 1999). Since dioecious species are typically perennials (Field *et al*., 2013a) and some can reproduce both clonally and sexually, the degree and frequency of flowering and clonal propagation can also both influence sex ratios. Several studies examining this in wild accessions of *Salix* have shown a female sex ratio bias (Barrett *et al*., 2010). Male bias is commonly seen in trees and often associated with biotic pollen dispersal in contrast to female-biased ratios observed in shrubs and herbs that tend to be clonal, perennial species (Field *et al*., 2013). Several studies of wild accessions of *Salix* have found sex ratio biases toward females (Ueno *et al*., 2007; Che-Castaldo *et al*., 2015), but male bias was demonstrated in a controlled cross experiment in *S. viminalis* (Alström-Rapaport *et al*., 1997). Some studies of experimental populations (Mosseler and Zsuffa, 1989) have revealed greater variability of sex ratio bias than in natural populations (Myers-Smith and Hik, 2012), but sex ratio bias in progeny of controlled crosses depends on the nature of the cross (inter- or intraspecific) as well as the ploidy levels of the parents.

The short generation time of *Salix* makes this genus suitable for studying sexual dimorphism and its genetic control. This study examined *S. purpurea* L. (purple osier willow), a naturalized species in North America and an important species in breeding shrub willow bioenergy crops in North America as it has been used in over 30% of all intra- and interspecific hybrids produced to date (Smart and Cameron, 2008). Critical traits to study for dimorphism include pest and disease resistance, drought tolerance, nitrogen and water use efficiency, and biomass yield.

This study describes the first investigation of sexual dimorphism in a *S. purpurea* diversity panel and F_1_ and F_2_ populations. The objectives were to (1) evaluate the phenotypic variability within naturally occurring and bred genotypes of *S. purpurea* (2) determine if there is sexual dimorphism of secondary sex characteristics among natural and bred populations and (3) test if observed sex ratios fit those expected based on a single locus sex determination system or if there is evidence for an multigenic control of sex ratio bias within the species.

## MATERIALS AND METHODS

### Germplasm and field trials

Three populations of *S. purpurea* L. were used in this study: a diversity panel of unrelated natural accessions, an F_1_ family, and an F_2_ family generated by crossing F_1_ individuals. The diversity panel contained 78 genotypes of *S. purpurea* natural accessions (Lin *et al*., 2009, Gouker *et al*., 2019) (Appendix S1). In July 2012, all genotypes were hand planted using 20-cm cuttings in a common garden design at three experimental sites (Appendix S2): Cornell AgriTech in Geneva, NY the Cornell Lake Erie Research and Extension Lab (CLEREL) in Portland, NY; and the West Virginia University (WVU) Agronomy Farm in Morgantown, WV. All sites were planted in a randomized complete block design with six replicates of four-plant plots at each location in single-row spacing with 1.82 m between rows and 0.40 m between plants within rows. Border rows containing either genotype 94006 or cultivar ‘Fish Creek’ were planted on the perimeter to avoid edge effects. At the end of the establishment year, all plants were coppiced and trials were measured in 2013 and 2014 and subsequently harvested and weighed in early 2015. Prior to re-growth of the second rotation in 2015, 112 kg ha^-1^ N-P-K fertilizer was applied to half of the replicates at each location to test for nitrogen utilization. Rust was surveyed at two locations at the end of the 2015 growing season.

The intraspecific F1 *S. purpurea* family was generated from a cross between the female genotype 94006 and the male genotype 94001, which were accessions collected near Syracuse, NY and were also present in the diversity panel. Two F_1_ siblings from this family were selected and crossed (‘Wolcott’ × ‘Fish Creek’) to generate the F_2_ population. A total of 100 F_1_ and 482 F_2_ progeny and their parents were hand planted using 20-cm cuttings at Cornell AgriTech in June 2014 (Appendix S2) in a randomized complete block design with four replicate blocks of three-plant plots in the same single-row spacing described above (Carlson *et al*., 2019). To avoid edge-effects, border rows containing 94006 and ‘Fish Creek’ were planted along the perimeter of the trial. At the end of the establishment year, all plants were coppiced, fertilized with 112 kg ha^-1^ N, 67 kg ha^-1^ P and K, and measurements were collected in 2015.

### Phenotyping

The diversity panel was evaluated for 26 biomass, morphological, phenological, physiological and wood composition traits measured as described (Appendix S3) across three sites in 2013 and 2014 and a subset of traits as well as rust incidence were measured in 2015.

The F_1_ and F_2_ populations were evaluated for all traits on one site in 2015 (Table 1) (Carlson *et al*., 2019). For growth measurements in the diversity panel, the inner two plants of each four-plant plot were measured and the central plant in each three-plant plot were measured in the F_1_ and F_2_ families. Rust was surveyed by visually assessing percent uredospore pustule coverage on leaves (0-100%) of all the plants in each plot. Sex was scored for clonally propagated plants of each genotype growing in nursery beds and was confirmed in experimental plots.

**Table 1.** Phenotypic traits measured in the *S. purpurea* F_1_ F_2_, and diversity panel.

| Trait | Abbreviation | Units |
| --- | --- | --- |
| ***Biomass*** |  |  |
| Height | HT | m |
| Stem number | STNo | # |
| Stem diameter | DIA | mm |
| Total stem area | SA | cm<sup>2</sup> |
| Internode length | INL | cm |
| Biomass Yield | YLD | dry Mg ha<sup>-1</sup> |
| ***Foliar*** |  |  |
| Leaf length | LFL | cm |
| Leaf width | LFW | cm |
| Leaf area | LFA | cm<sup>2</sup> |
| Leaf perimeter | LFP | cm |
| Leaf dry weight | LFDW | g |
| Specific leaf area | SLA | cm<sup>2</sup> g<sup>-1</sup> |
| ***Architecture*** |  |  |
| Crown diameter | CDIA | cm |
| Crown form | FORM | degrees ° |
| ***Composition*** |  |  |
| Hemicellulose | HCL | % |
| Cellulose | CLS | % |
| Lignin | LIG | % |
| Ash | ASH | % |
| Specific gravity | SPGR | g cm<sup>-3</sup> |
| ***Physiology*** |  |  |
| August SPAD | SPAD1 | SPAD units |
| September SPAD | SPAD2 | SPAD units |
| Stomatal<br>conductance | $g_s$ | mmol m<sup>-2</sup> s<sup>-1</sup> |
| Canopy color | RGB | RGB |
| Survival | SRV | % |
| ***Phenology*** |  |  |
| Vegetative<br>phenology | VPHE | day of year |
| Floral phenology | FPHE | day of year |
| ***Pathology*** |  |  |
| Rust severity | RUST | % |

### Statistical analysis

Statistical analyses were conducted using SAS^®^ version 9.4 (SAS Institute, Cary, NC) and R version 3.2.3 (R Core Development Team). Mixed linear models were used to analyze phenotypic data implemented in SAS with the PROC MIXED statement and using the *lmer* function within the *lme4* package in R (Bates *et al*., 2015). All dependent variables were tested for homogeneity of variances and normality using PROC UNIVARIATE using Kolmogorov-Smirnov D and Shapiro-Wilk’s K statistics. Non-parametric methods were used when phenotypes that were not normally distributed could not be transformed to meet the assumptions of parametric analyses using Box-Cox powers or log-transformation. Yield data were square-root transformed to meet assumptions of normality. Pearson’s product-moment correlations (*r*) were calculated among all traits. To test for statistical differences between phenotypic traits based on sex, a two-tailed Mann-Whitney *U*test was conducted with hermaphrodite genotypes excluded. To estimate the predictability and relationship of each trait and biomass yield, multiple linear regression was performed with PROC REG using the stepwise regression method. Model adequacy was checked with a general linear model to assess the global significance with PROC GLM.

## RESULTS

### Phenotypic variation

In the diversity panel, all traits showed large differences among genotypes (Appendix S4). Effects of genotype, location, and genotype × location were highly significant for yield (*P*<0.05) (Table 2). Overall, the greatest differences in growth traits between genotypes were for stem area (SA) and stem height (HT) (Fig. 1). The range of values across two growing seasons in the diversity panel were 0.13 to 84.38 cm^2^ for SA and 0.11 to 4.88 m in HT (Appendix S4). There was also wide variability in stem number (STNo), which increased on average by 22% from the first to the second year. Crown form (FORM), calculated from crown diameter (CDIA), ranged from approximately 9 to 88° mean branching angle, but all genotypes had variable FORM across sites. Of the four metrics obtained from leaf scans, the greatest variation was for leaf perimeter (LFP). The same degree of variability was observed across sites for phenology and physiology traits, where stomatal conductance (*g_s_*) had the greatest variability with the maximum value of 1164.2 mmol m^-2^ s^-1^ and the minimum value of 45.5 mmol m^-2^ s^-1^ in year 1 (Appendices S5, S6). The genotypic means for wood composition and specific gravity (SPGR) also had wide variances. The largest variation was observed for cellulose content (CLS) with a range of 37.94% difference between the highest and lowest value. On average, second year measurements of hemicellulose content (HCL) and CLS were greater than first year values by 0.79% and 10.68%, respectively (Appendix S4). Lignin content (LIG) decreased by 5.48% in year 2 compared to year 1 and ash content (ASH) declined by 27.76%, but SPGR only decreased by 0.04 g cm^-1^ in the second year.

**Table 2.** Mixed model test of effects of genotype and location on yield.

| **Source** | **DF** | ***F* Ratio** | **Pr > *F*** |
| --- | --- | --- | --- |
| Location | 2 | 155.23 | <0.0001* |
| Genotype | 77 | 11.51 | <0.0001* |
| Genotype × Location | 154 | 1.80 | <0.0001* |

\*Significant differences at $P < 0.05$

**Figure 1.**
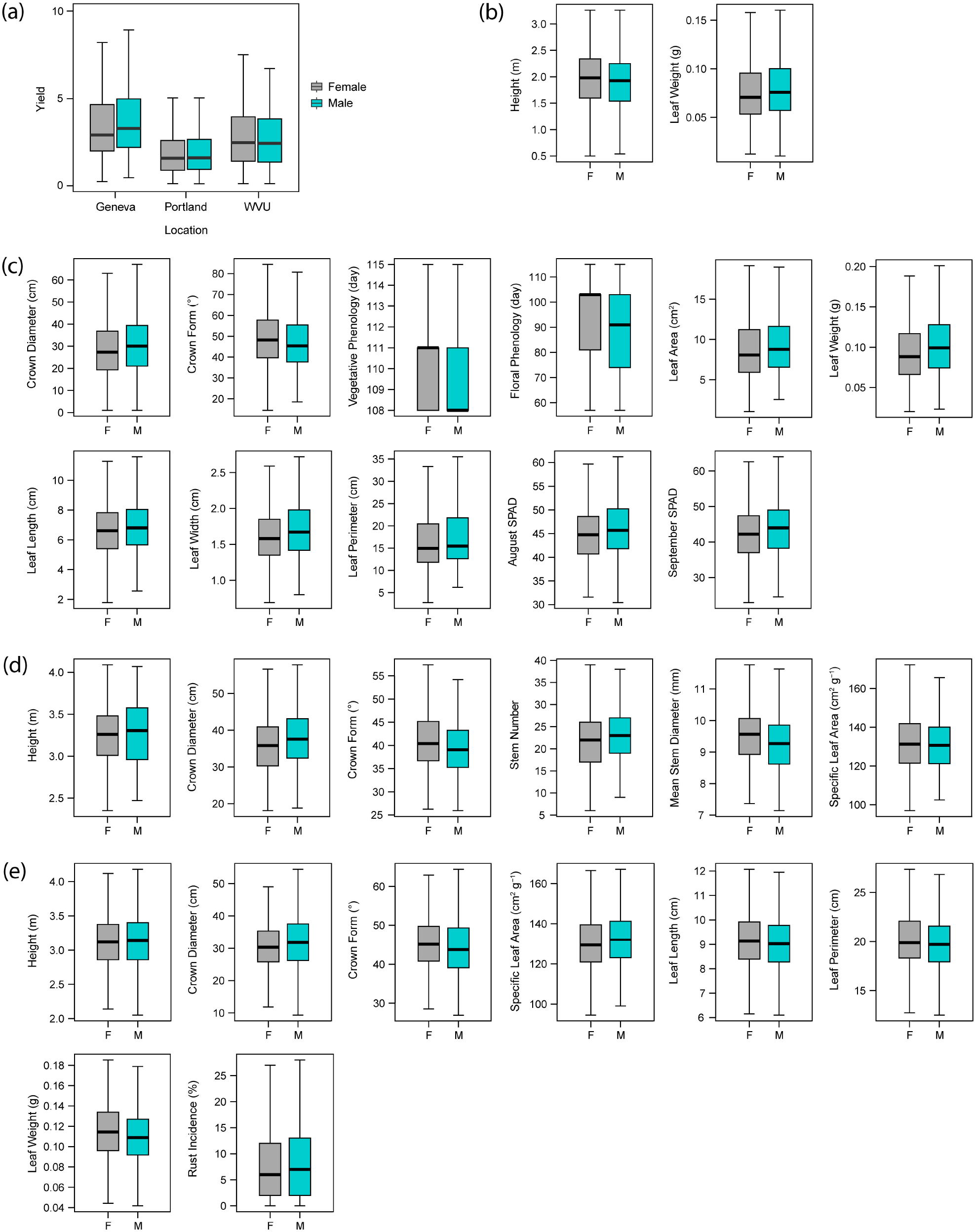
Box plots of biomass and morphological traits that were significantly different (*P*<0.05) between females (grey boxes) and males (blue boxes). (A) Second year, first rotation biomass yield from the diversity panel measured across three field trials. (B) First year, significantly different traits from the diversity panel. (C) Second year, significantly different traits from the diversity panel. (D) Significantly different traits from F1 population. (E) Significantly different traits from F_2_ population.

In the F_1_ and F_2_ families, mean growth in Geneva was better than that of the diversity panel (Table 3, Appendix S7, Appendix S8). The means for SA and HT for first-year coppice growth were greater in the F_1_ and F_2_ families compared to the diversity panel. The first-year mean SA in the diversity panel in Geneva was 14.95 cm^2^, while it was 16.87 and 12.62 cm^2^ in the F_1_ and F_2_ families, respectively. Relative differences in first-year post-coppice HT were similar, as the diversity panel in Geneva had mean HT of 2.30 m, the mean HT in the F_1_ family was 3.25 m and the mean HT of the F_2_ family was 3.11 m. SPAD measurements and specific leaf area (SLA) showed similar trends with the diversity panel with lower means for both traits compared to the F_1_ and F_2_ families (Fig. 2).

**Table 3.** Means and standard deviations of phenotypic traits in the *S. purpurea* intraspecific F_1_ family (n=100) and F_2_ family (n=482) in Geneva, NY.

|  | F<sub>1</sub> *S. purpurea* family |  | F<sub>2</sub> *S. purpurea* family |  |  |
| --- | --- | --- | --- | --- | --- |
| Trait<sup>1</sup> | Mean ±SE | Min - Max | Mean ±SE | Min - Max | *t*-value<sup>2</sup> |
| ***Biomass*** |  |  |  |  |  |
| HT | 3.26 ±0.19 | 2.15 - 4.78 | 3.11 ±0.09 | 0.98 - 4.31 | 14.7* |
| STNo | 21.8 ±0.32 | 6.00 - 41.0 | 18.7 ±0.16 | 1.00 - 44.0 | 8.51* |
| SDIA | 9.44 ±0.05 | 6.36 - 12.4 | 8.81 ±0.02 | 5.00 - 12.4 | 12.9* |
| SA | 16.9 ±0.28 | 4.48 - 33.3 | 12.6 ±0.12 | 2.80 - 37.7 | 7.30* |
| ***Foliar*** |  |  |  |  |  |
| LFL | 9.84 ±0.06 | 6.52 - 13.9 | 9.15 ±0.03 | 4.58 - 18.9 | 9.97* |
| LFW | 2.18 ±0.02 | 1.40 - 4.93 | 2.04 ±0.01 | 1.10 - 12.7 | 4.90* |
| LFA | 17.2 ±0.19 | 8.19 - 33.8 | 14.9 ±0.08 | 4.08 - 37.9 | 11.1* |
| LFP | 22.4 ±0.28 | 13.8 - 57.8 | 21.0 ±0.12 | 9.54 - 58.2 | 4.80* |
| LFDW | 0.13 ±0.002 | 0.06 - 0.28 | 0.11 ±0.001 | 0.03 - 0.41 | 9.74* |
| SLA | 134 ±0.89 | 97.0 - 233 | 132 ±0.42 | 61.9 - 361 | 1.15 |
| ***Architecture*** |  |  |  |  |  |
| CDIA | 36.9 ±0.43 | 18.1 - 84.7 | 31.3 ±0.19 | 3.10 - 77.0 | 12.6* |
| FORM | 40.4 ±0.32 | 19.8 - 59.3 | 45.2 ±0.17 | 21.6 - 84.2 | 12.1* |
| ***Physiology*** |  |  |  |  |  |
| SPAD1 | 55.9 ±0.29 | 11.1 - 76.3 | 57.2 ±0.13 | 26.2-91.9 | -3.97* |
| RGB | 112 ±0.78 | 55.5 - 158 | 111 ±0.37 | 48.3-168 | 0.90 |
| SRV | 99.7 ±0.20 | 33.3 - 100 | 99.1 ±0.15 | 0.00 - 100 | 2.70* |
| ***Pathology*** |  |  |  |  |  |
| RUST | - | - | 0.08 ±0.002 | 0.00-0.86 | - |

<sup>1</sup>Phenotypic traits were measured in 2015. See Materials and Methods for trait definitions and Table 1 for abbreviations
<sup>2</sup>Student's *t*-test statistic, where \* denotes significant differences among populations at a *P*<0.01 level-of-confidence, with positive values indicating greater means in the F<sub>1</sub> and negative value indicating greater means in the F<sub>2</sub>.

**Figure 2.**
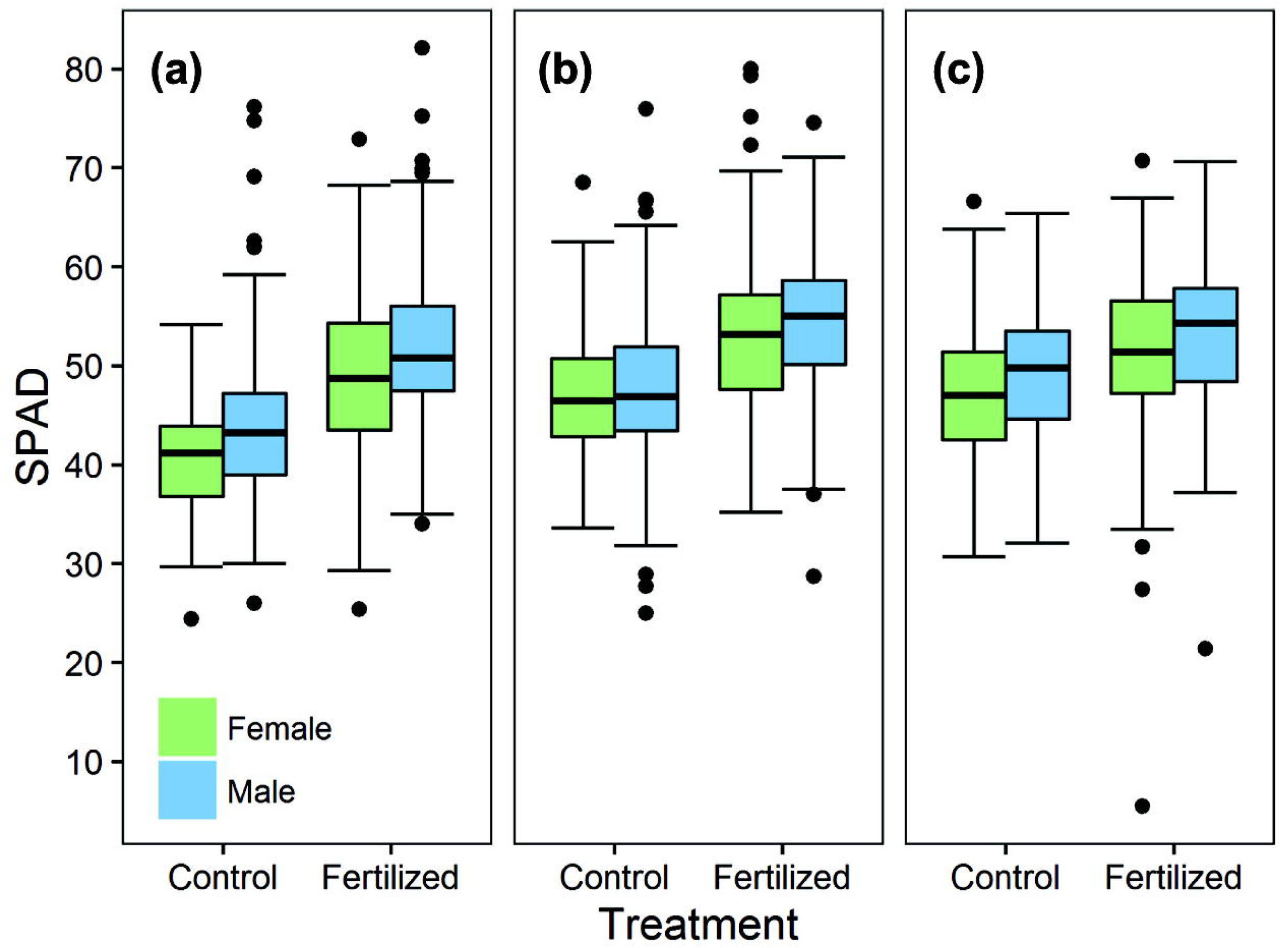
SPAD values for monitoring nitrogen utilization in *Salix purpurea* diversity panel. Box plots representing females are colored in green and males colored in blue. SPAD values for control and fertilized plots for (A) Geneva, NY (F_1_,_105_ = 15.73, *P* < 0.01), (B) Portland, NY (F1,105 = 3.44, *P* = 0.06), and (C) Morgantown, WV (F1,105 = 5.96, *P* = 0.02).

In general, overall lower trait means were observed for biomass and physiological traits in the F_2_ family compared to the F_1_ family (Table 3). For instance, SDIA, HT, STNo, and SA were all significantly greater (*P* < 0.01) in the F_1_ family with *t*-values ranging from 7.3 to 14.7, as well as traits related to stem architecture, CDIA (*t* = 12.6) and FORM (*t* = 12.1) (Table 3). Stem area was ~33% greater in the F_1_ family compared to the F_2_ family in 2015. Although morphological leaf traits were significantly greater in the F_1_ family, SLA and canopy color (RGB) were the only two traits that were not significantly different between F_1_ and F_2_ families, whereas SPAD was the only trait that was significantly greater in the F_2_ family (*t* = −3.97, *P* < 0.01).

### Sexually dimorphic phenotypes

Significant sex differences were found in *S. purpurea*, with males producing greater growth and significantly greater means for most traits measured (Fig. 1, Table 4, Table 5). For the diversity panel in the first year of growth (2013), two traits were significantly dimorphic (*P*<0.05, Table 4). Mean HT of females (1.95 m) across all three sites was 3.7% greater than males (1.88 m). Yet males had significantly heavier leaf dry weight (LFDW) than females. The number of dimorphic traits increased in year two (2014), when 11 of the traits were significantly dimorphic (Table 4). There were greater trait means in males for CDIA, leaf length (LFL), leaf width (LFW), leaf area (LFA), LFP, and LFDW. Crown form (FORM) was calculated from CDIA and showed significantly lower branching angle in the males reflecting a greater crown diameter in males than in females. Mean floral (FPHE) and vegetative (VPHE) phenology measurements showed greater means for females indicating earlier bud break for males (Table 4). Six traits were sexually dimorphic in the F_1_ family (Table 5). Male means for HT and specific leaf area (SLA) were greater than for females, as was CDIA, meaning that the FORM angle was lower in males than in females. In the F_1_ family, SDIA of female progeny was greater than that of males, whereas STNo was greater in males compared to females. In the F_2_ family, HT was significantly greater for males, as it was in the F_1_ family. LFDW, LFP, and LFL means were greater in females than males, while CDIA was greater for males, with a lower form angle, as was observed in the diversity panel and F1 family.

**Table 4.** Comparison of phenotypic traits for female and male individuals in the *S. purpurea* diversity panel across three growing seasons.

| Trait | Female (n=49) |  | Male (n=29) |  | P | Diff (%) |
| --- | --- | --- | --- | --- | --- | --- |
|  | Mean ± SE | CV (%) | Mean ± SE | CV (%) |  |  |
| 2013 |  |  |  |  |  |  |
| HT | 1.95 ±0.02 | 28.04 | 1.88 ±0.02 | 28.81 | 0.05** | -3.29 |
| STNo | 18.14 ±0.34 | 54.76 | 17.63 ±0.45 | 58.43 | 0.35 | -2.82 |
| SDIA | 7.3 ±0.06 | 22.62 | 7.38 ±0.07 | 21.62 | 0.38 | 1.22 |
| SA | 9.4 ±0.24 | 75.84 | 9.14 ±0.31 | 76.82 | 0.49 | -2.75 |
| INL | 13.58 ±0.17 | 37.74 | 13.38 ±0.22 | 37.19 | 0.55 | -1.47 |
| LFL | 6.91 ±0.07 | 23.58 | 7.02 ±0.09 | 23.68 | 0.35 | 1.62 |
| LFW | 1.69 ±0.05 | 72.05 | 1.85 ±0.07 | 67.64 | 0.06 | 9.31 |
| LFA | 8.86 ±0.18 | 49.53 | 9.42 ±0.24 | 47.89 | 0.08* | 6.32 |
| LFP | 23.9 ±1 | 101.00 | 24.19 ±1.33 | 102.54 | 0.85 | 1.18 |
| LFDW | 0.0891 ±0.001 | 42.76 | 0.0896 ±0.003 | 58.78 | 0.01*** | 9.17 |
| SLA | 128.14 ±3.73 | 70.66 | 131.54 ±5.33 | 75.42 | 0.56 | 2.65 |
| HCL | 17.54 ±0.04 | 5.26 | 17.67 ±0.05 | 5.43 | 0.10* | 0.72 |
| CLS | 37.55 ±0.13 | 8.53 | 37.55 ±0.18 | 9.14 | 0.98 | 0.02 |
| LIG | 28.9 ±0.09 | 7.43 | 28.89 ±0.11 | 7.37 | 0.96 | -0.03 |
| ASH | 2.16 ±0.02 | 26.87 | 2.18 ±0.03 | 28.82 | 0.44 | 1.23 |
| SPGR | 0.45 ±0 | 15.97 | 0.45 ±0.004 | 14.69 | 0.96 | 0.01 |
| SPAD1 | 45.85 ±0.32 | 20.50 | 45.88 ±0.42 | 20.69 | 0.94 | 0.08 |
| SPAD2 | 41.2 ±0.35 | 20.34 | 42.29 ±0.53 | 23.36 | 0.06* | 2.67 |
| g<sub>s</sub> | 596.39 ±6.99 | 32.84 | 591.92 ±8.63 | 31.41 | 0.67 | -0.75 |
| 2014 |  |  |  |  |  |  |
| HT | 3.16 ±0.02 | 21.62 | 3.1 ±0.03 | 22.51 | 0.25 | -1.84 |
| STNo | 22.14 ±0.39 | 52.00 | 21.55 ±0.53 | 56.01 | 0.39 | -2.67 |
| SDIA | 239.42 ±5.13 | 63.58 | 172.35 ±4.69 | 62.12 | 0.07* | -28.01 |
| SA | 20.96 ±0.47 | 66.45 | 21.19 ±0.62 | 66.52 | 0.78 | 1.07 |
| ILEN | 13.59 ±0.24 | 42.07 | 12.98 ±0.28 | 40.60 | 0.05** | -4.45 |
| YLD | 2.71 ±0.06 | 69.25 | 2.79 ±0.08 | 67.81 | 0.56 | 2.81 |
| LFL | 6.6 ±0.06 | 28.18 | 6.92 ±0.07 | 24.54 | <0.01*** | 4.88 |
| LFW | 1.81 ±0.06 | 96.01 | 2.03 ±0.11 | 118.12 | 0.05** | 12.22 |
| LFA | 8.72 ±0.14 | 46.06 | 9.49 ±0.17 | 41.80 | <0.01*** | 8.76 |
| LFP | 17.51 ±0.3 | 51.03 | 18.38 ±0.37 | 45.54 | 0.04** | 4.97 |
| LFDW | 0.09 ±0.001 | 40.16 | 0.11 ±0.002 | 40.04 | <0.01*** | 12.17 |
| SLA | 92.28 ±0.75 | 24.10 | 90.56 ±0.75 | 19.03 | 0.15 | -1.86 |
| CDIA | 29.09 ±0.5 | 50.60 | 31.42 ±0.74 | 53.59 | 0.01*** | 8.01 |
| FORM | 49.36 ±0.46 | 27.69 | 47.61 ±0.64 | 30.84 | 0.03** | -3.55 |
| HCL | 17.69 ±0.03 | 4.36 | 17.79 ±0.04 | 4.34 | 0.09* | 0.55 |
| CLS | 41.61 ±0.07 | 4.22 | 41.48 ±0.09 | 4.26 | 0.33 | -0.32 |
| LIG | 27.29 ±0.05 | 4.42 | 27.36 ±0.06 | 4.37 | 0.42 | 0.26 |
| ASH | 1.55 ±0.02 | 24.89 | 1.58 ±0.02 | 25.22 | 0.34 | 1.84 |
| SPGR | 0.49 ±0.002 | 9.91 | 0.49 ±0.002 | 7.82 | 0.85 | -0.14 |
| **SPAD1** | **44.93 ±0.24** | **12.84** | **46.15 ±0.33** | **13.29** | **<0.01\*\*\*** | **2.73** |
| **SPAD2** | **42.45 ±0.28** | **18.35** | **43.72 ±0.36** | **17.66** | **<0.01\*\*\*** | **2.99** |
| $g_s$ | 474.51 ±5.7 | 35.66 | 483.68 ±7.18 | 33.88 | 0.39 | 1.93 |
| **VPHE** | **110.54 ±0.14** | **3.12** | **108.88 ±0.21** | **3.64** | **<0.01\*\*\*** | **-1.51** |
| **FPHE** | **95.25 ±0.55** | **13.95** | **86.98 ±0.85** | **18.25** | **<0.01\*\*\*** | **-8.69** |
| **2015** |  |  |  |  |  |  |
| **RUST** | **28.33±0.53** | **43.70** | **25.45±0.72** | **50.49** | **0.01\*\*\*** | **-10.17** |

All values are mean ± SE across three locations except for RUST which was evaluated across two locations.
Two-tailed Mann-Whitney $U$ -test (df=1) results, where significant values ( $P<0.05$ ) are denoted by bold font.
Additional levels significance are also shown denoted by $P<0.1$ (\*), $P<0.05$ (\*\*) and bold font, $P<0.01$ (\*\*\*)
Positive values for dimorphism denote male-biased difference and negative values denote female-biased difference.

**Table 5.** Comparison of phenotypic traits for male and female individuals in an intraspecific F_1_ *S. purpurea* family (n=100) and a F_2_ *S. purpurea* family (n=482) measured in 2015 in Geneva, NY.

| Trait | F<sub>1</sub> *S. purpurea* family |  |  |  |  |  | F<sub>2</sub> *S. purpurea* family |  |  |  |  |  |
| --- | --- | --- | --- | --- | --- | --- | --- | --- | --- | --- | --- | --- |
|  | Female (n=70) |  | Male (n=30) |  | P | Diff (%) | Female (n=266) |  | Male (n=216) |  | P | Diff (%) |
|  | Mean ±SE | CV | Mean ±SE | CV |  |  | Mean ±SE | CV | Mean ±SE | CV |  |  |
| HT | **3.24 ±0.22** | **11.42** | **3.29 ±0.34** | **10.94** | **0.02** | **1.54** | **3.11 ±1.15** | **12.22** | **3.13 ±1.35** | **12.78** | **0.02** | **0.64** |
| STNo | **21.2 ±0.4** | **33.33** | **23.3 ±0.53** | **26.09** | **0.03** | **9.52** | 18.4 ±0.21 | 38.89 | 19.0 ±0.25 | 36.84 | 0.11 | 5.56 |
| SDIA | **9.52 ±0.06** | **9.87** | **9.26 ±0.09** | **10.91** | **<0.01** | **-2.73** | 8.82 ±0.03 | 9.75 | 8.81 ±0.03 | 10.67 | 0.43 | -0.11 |
| SA | 16.8 ±0.34 | 33.81 | 17.1 ±0.49 | 31.05 | 0.22 | 1.79 | 12.4 ±0.16 | 41.69 | 12.9 ±0.19 | 43.64 | 0.12 | 4.03 |
| LFL | 9.84 ±0.07 | 11.99 | 9.83 ±0.12 | 12.82 | 0.85 | -0.10 | **9.19 ±0.04** | **13.38** | **9.09 ±0.04** | **14.52** | **0.03** | **-1.09** |
| LFW | 2.19 ±0.02 | 15.98 | 2.16 ±0.04 | 18.98 | 0.32 | -1.37 | 2.05 ±0.01 | 21.95 | 2.04 ±0.02 | 27.94 | 0.13 | -0.49 |
| LFA | 17.2 ±0.23 | 22.33 | 17.1 ±0.4 | 25.26 | 0.56 | -0.58 | 15.1 ±0.11 | 23.71 | 14.8 ±0.12 | 24.73 | 0.07 | -1.99 |
| LFP | 22.2 ±0.33 | 24.86 | 22.7 ±0.53 | 25.07 | 0.51 | 2.25 | **21.1 ±0.16** | **25.21** | **20.8 ±0.18** | **26.11** | **0.02** | **-1.42** |
| LFDW | 0.13±0.002 | 23.08 | 0.13±0.003 | 30.77 | 0.82 | 0.00 | **0.12 ±0.000** | **25.00** | **0.11 ±0.001** | **27.27** | **<0.01** | **-8.33** |
| SLA | 133 ±1.02 | 12.84 | 132 ±1.58 | 12.82 | 0.58 | -0.75 | **131 ±0.56** | **13.89** | **134 ±0.59** | **13.04** | **<0.01** | **2.26** |
| CDIA | **36.3 ±0.49** | **22.59** | **38.7 ±0.88** | **24.44** | **0.02** | **6.61** | **30.6 ±0.24** | **25.23** | **32.1 ±0.3** | **27.32** | **<0.01** | **4.90** |
| FORM | **40.9 ±0.38** | **15.45** | **39.1 ±0.6** | **16.47** | **0.02** | **-4.40** | **45.8 ±0.22** | **16.05** | **44.6 ±0.27** | **17.96** | **<0.01** | **-2.62** |
| SPAD1 | 56.2 ±0.3 | 8.52 | 54.9 ±0.68 | 12.28 | 0.11 | -2.31 | 57.2 ±0.19 | 10.05 | 57.1 ±0.19 | 8.88 | 0.88 | -0.17 |
| RGB | 112 ±0.94 | 14.11 | 111 ±1.42 | 13.78 | 0.38 | -0.89 | 111 ±0.49 | 14.50 | 112 ±0.55 | 14.46 | 0.08 | 0.90 |
| SRV | 99.6 ±0.26 | 4.44 | 99.7 ±0.29 | 3.10 | 0.87 | 0.10 | 99.34 ±0.17 | 5.51 | 98.8 ±0.25 | 7.43 | 0.05 | -0.54 |
| RUST | - | - | - | - | - | - | **7.9 ±0.002** | **88.61** | **8.8 ±0.003** | **90.91** | **0.03** | **11.39** |

Values are mean ± SE.
Two-tailed Mann-Whitney *U*-test (d.f.=1) results, where significant values ( $P<0.05$ ) are denoted by bold font.
Positive values for dimorphism denote male-biased difference and negative values denote female-biased difference.

**Table 6.** Mixed model test for nitrogen utilization.

| **Source** | **DF** | **F Ratio** | **Pr > F** |
| --- | --- | --- | --- |
| Location | 2 | 65.26 | < **0.0001** * |
| Treatment | 1 | 170.00 | < **0.0001** * |
| Sex | 1 | 15.75 | **0.0001** * |
| Sex × Treatment | 1 | 0.31 | 0.58 |
| Location × Treatment | 2 | 6.51 | < **0.01** * |

\*Significantly different at $P < 0.05$

Leaf rust severity (RUST) was surveyed during the 2015 growing season in the diversity panel and F_2_ family. In the diversity panel in 2015, there were significant differences between the two locations surveyed (*P*<0.05). Based on percent disease severity using least square means, for Geneva, NY, females had a greater mean score (25%) for RUST than males (22%) and in Portland, NY least square means for RUST were 30% and 27% for females and males, respectively, with overall significant differences of RUST averaged across locations between males and females (Table 4, Fig. 3A). The male parents of the F_1_ and F_2_ families had significantly greater mean RUST scores than the female parents (Fig. 3B). Similarly, the male F_2_ progeny had significantly greater mean RUST than the F_2_ female progeny (*P*=0.02) (Table 5, Fig. 3B). The overall F_2_ progeny means for RUST were greater than that of the female parent, ‘Wolcott’, but less than that of the male parent ‘Fish Creek’. Overall, there was a significant negative correlation between RUST and both SA and HT (*P*<0.05), with a significant positive correlation between RUST infection and SPAD measurements (*P*<0.01) (Appendix S9).

**Figure 3.**
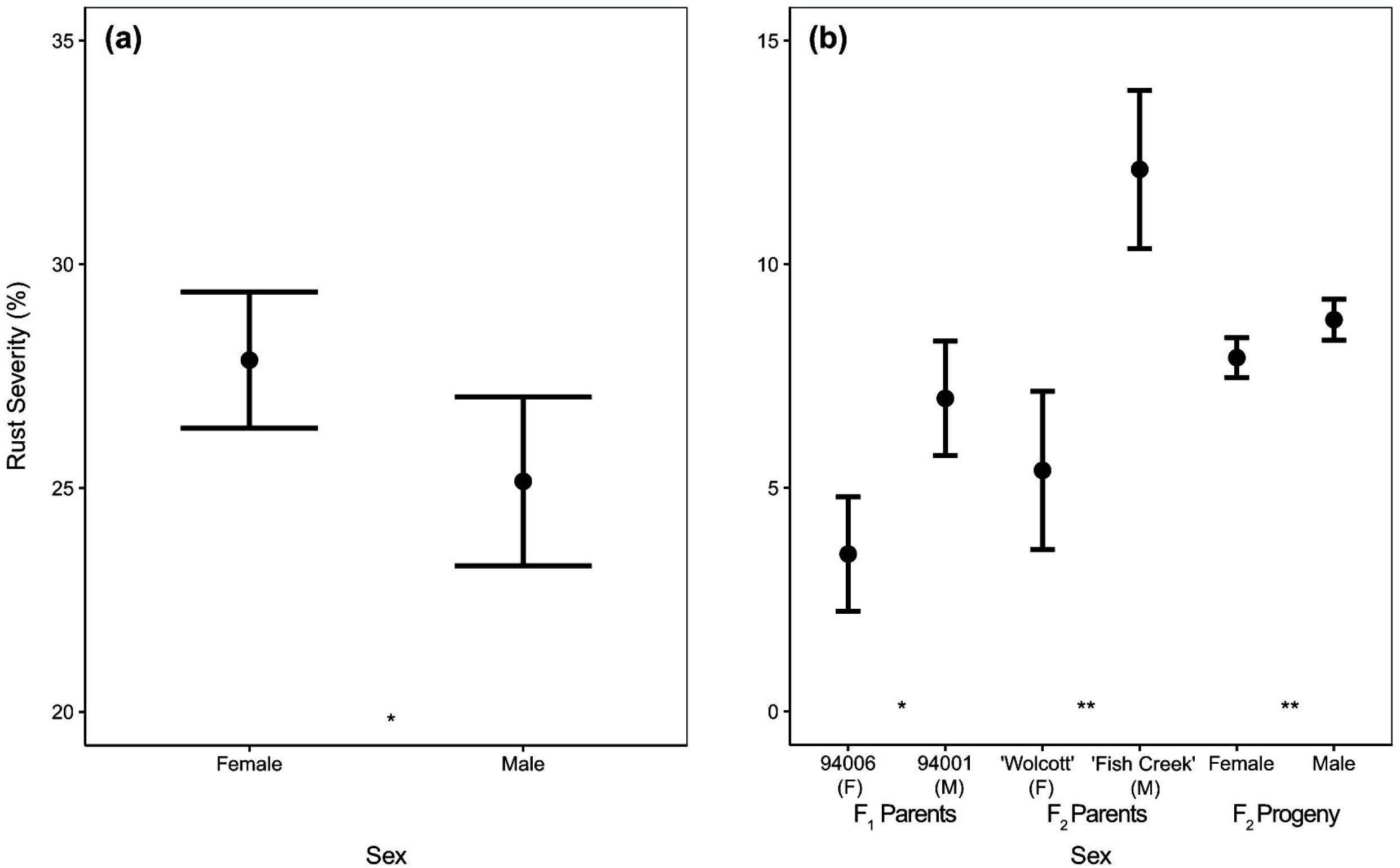
Least square means for leaf rust severity scores of female and male *Salix purpurea*. (A) Rust severity scores and standard errors on females and males of the diversity panel at the Geneva, NY and Portland, NY field sites. (B) Rust severity scores and standard error on F_1_, F_2_ parents and female and male F_2_ progeny in Geneva, NY. Significant differences between females and males within each site and population are denoted by * *P* < 0.10, and ** *P* < 0.01, where n.s. denotes no significant difference.

### Sex ratios

There were 49 females and 29 males in the diversity panel, which were confirmed across years and experimental locations based on documented sex phenotypes in nursey beds. The diversity panel was not necessarily representative of the sex ratios in natural populations, since it was assembled from only a few selected individuals from multiple sites, rather than a thorough sampling of all individuals of a population (Gouker, *et al*., 2019). There was significant departure from the expected 1: 1 segregation ratio of males and females in both the F1 and F_2_ families. The F_1_ family consisted of 70 females and 30 males (ratio=2.33:1, *P* <0.01) and the F_2_ family contained 266 females and 216 male genotypes (ratio=1.23:1, *P* =0.02).

### Allometric model for yield

All measured traits from the diversity panels were used as parameters in allometric models to identify relationships between YLD and the yearly growth measurements using multiple linear regression to predict second year biomass. Separating genotypes by sex and examining allometric relationships with YLD revealed no significant difference (*P*=0.15) or advantage of predicting YLD and therefore the data were not distinguished by sex in the regression model. Variable inflation factors greater than 10 were observed between total SDIA and total SA and indicated multicollinearity as indicated with the high correlation coefficient of r=0.95 (Appendix S10). Since SA had a greater correlation with YLD and explained a greater percentage of the variance in the model, it was kept and SDIA was removed. All other variables that did not meet the *P*<0.05 significance level were also removed. To test for global significance of variables, a general linear model was fitted and revealed SLA in 2014 (*P*=0.88) and LFP (*P*=0.84) were insignificant and were also removed as predictor variables. The best predictors for the final multiple linear regression model were SA in 2013 and 2014, HT in 2014, and AugSPAD in 2014 (Appendix S5), where yearly SA measurements gave the most accurate estimates of YLD. A strong positive fit of predicted and observed biomass YLD resulted in an overall R^2^= 0.79 (Appendix S10).

## DISCUSSION

For dioecious species, genetic factors, selection pressures over time, and ecological adaptation can lead to differential fitness between males and females and result in dimorphism of secondary characteristics (Sakai and Weller, 1999). While such differential selection in males and females may be expected to lead to sex ratio bias, very few published records of such bias exist.

Sex ratio bias has been reported in *Salix* spp. and is most often biased towards females in an approximate 2 female:1 male ratio (Alström-Rapaport *et al*., 1997; Rottenberg, 1998; Dudley, 2006; Ueno *et al*., 2007; Hughes *et al*., 2009; Myers-Smith and Hik, 2012), though the closest taxonomically related genus, *Populus*, has shown male-biased ratios (Tuskan *et al*. 2012). The female bias observed in the F_1_ and F_2_ families of this study may be explained by the occurrence of pollen competition (certation). Especially when pollen load is high, as in the case with controlled crosses, the female determining pollen may be inherently more successful at fertilization. We can speculate that there was relaxation of female:male ratios from the F_1_ to F_2_ generation as a result of slight inbreeding depression, differential mortality, or environmental factors.

Despite the female-biased sex ratios in the F_1_ and F_2_ families, our results showed that males were superior to females for many traits. Comparison of coefficients of variation (CV) between sexes in each population showed consistently greater variation in males, which may indicate that males have greater plasticity in response to environmental conditions, or alternatively that females spend more resources on seed production and cannot vary growth responses in relation to different environmental conditions. Yield and traits positively correlated with yield also showed greater trait means in males. Direct measurement of YLD was not significantly different, however. Within the F_1_ family, all dimorphic traits except for DIA were male-biased. It was shown that males have earlier bud break, which would extend their growing season and may partly explain differences in leaf traits (Tharakan *et al*., 2008). However, it has been suggested that other phenological events (i.e. leaf unfolding and duration, growth cessation and leaf abscission) affect annual biomass production, where growth cessation and late season leaf retention may also impact aboveground growth as well as nutrient recycling and storage *in planta* (Weih, 2009). An intensive study monitoring these additional traits could provide clues as to whether sex-specific physiological patterns are seen across other dioecious species as well.

Sex dimorphism has also been studied extensively in the closely-related genus *Populus*. Examination of phenotypic and gene expression data in *P. tremula* has shown no evidence of sexual dimorphism for morphological or biochemical traits (Robinson *et al*., 2014). Minor differences across a set of growth traits observed in *P. euphratica* suggested that male trees are more vigorous than females (Petzold *et al*., 2012). Studies of *P. deltoides* and *P. tremuloides* hybrids revealed significantly greater biomass production in males (Pauley, 1948; Farmer, 1964). When examining the evidence for sexual dimorphism in *Salix*, there are also opposing observations. In *S. planifolia*, it was reported that a larger allocation of resources is needed for reproduction in females than in males (Turcotte and Houle, 2001), which may lead to the assumption that females exhibit less biomass growth compared to males. Other studies have shown that females have growth rates similar and sometimes greater, but not significantly different, than males (Åhman, 1997; Sakai *et al*., 2006).

Data from this study revealed consistent trends of dimorphism for CDIA and the associated FORM showed a significant male bias for greater CDIA and subsequent shallower branching angle for all years and populations. Specific gravity contributes to the mechanical properties of wood and is known to scale positively with biomechanical strength and therefore directly influences plant architecture (Chave *et al*., 2009). This has practical importance for shrub willow, because plants with a wide CDIA that expands into the alleys of a field will result in biomass that may not be collected by a harvester. If specific cultivars have wide branching angles, this will result in a loss of harvestable biomass and reduction in yield.

Another interesting result of sexual dimorphism observed in this study was the significantly greater rust severity on male plants in the F_2_ family and the male progenitors, whereas in the diversity panel females exhibited greater rust severity. *Melampsora* leaf rust is the most severe plant disease affecting short-rotation willow plantations where long-term stability of yield will depend on host resistance. Resistance to *Melampsora* spp. has been mapped to multiple linkage groups in backcross and full-sib families in *S. viminalis* (Rönnberg-Wästljung *et al*., 2008; Hanley *et al*., 2011; Samils *et al*., 2011) with significant QTL for rust resistance on Chr01, Chr05, and Chr10 in *S. purpurea* (Carlson *et al*., 2019). Despite the sex dimorphism for RUST from this study in the F_2_ family, no resistance QTL have mapped to Chr15 near the SDR. Among the studies of rust severity in willow, there are few that have provided information on sex dimorphism. A study of largely unrelated commercial cultivars has indicated that there is greater rust severity on female plants (Moritz *et al*., 2016). Conversely, it has been observed in two other *Salix* studies that male-biased rust severity also exists (McCracken and Dawson, 2003; Pei *et al*., 2008). Our results also present conflicting evidence between the diversity panel and F_2_ family. The initial selection and cross of 94006 and 94001 had significantly different rust susceptibilities, as well as the full-sib F1 progeny ‘Wolcott’ and ‘Fish Creek’ which had greater rust severity than the parental genotypes. The subsequent cross of these full-sibs was used to generate the F_2_ full-sib family which also showed significantly greater rust susceptibility in male genotypes. Therefore, these differences may be a result of a genetic bottle neck through selective breeding which skewed rust susceptibility towards male genotypes.

Differences in observed severity of males and females between the diversity panel and the F_2_ family could be due to the timing of rust assessment. This may be particularly important for the higher level of rust incidence in Portland, NY, which was surveyed late in the season and disease had already advanced causing extensive pre-mature defoliation and rendering phenotypic differences that may exist between sexes indistinguishable. This suggests that differential rust susceptibility could also be driven by phenological differences between male and female plants. It is hypothesized that through life-history trade-offs of sexual morphs in dioecious species, females typically allocate greater resources towards reproduction and defense against pests and diseases (Seger and Eckhart, 1996; Vega-Frutis *et al*., 2013), and males invest more resources into primary growth (Delph, 1999; Obeso, 2002). Additionally, there may be differences in mechanical or biochemical defense mechanisms that we did not measure and for which there are limited studies examining this topic (Bañuelos *et al*., 2004).

The SPAD values observed in this study, used as a non-destructive method to quantify nitrogen status in the plant, showed significantly greater values in nitrogen amended versus control plots, but also significantly greater values in males than females in the diversity panel. The male-biased nitrogen capacity could possibly have been selected for by a greater nitrogen requirement for pollen production (Carolyn and Rundel, 1979, Boecklen *et al*., 1990) and the greater resource allocation towards primary growth. It may also be that females require a larger investment of nitrogen for seed production leaving more nitrogen available in males as observed by the SPAD readings. Although, a review conducted by Hultine (2016) examining differential resource acquisition between sexes of 22 species across multiple environments, concluded that females generally do not have greater nutrient uptake or efficiency compared with males under optimal growing conditions. Based on evidence so far, there tends to be greater nitrogen content in males plants in willow and other plant species, however, it is uncertain of the exact mechanisms contributing to these sexually dimorphic observations.

## CONCLUSIONS

Shrub willows include very diverse species used in a number of horticultural applications ranging from biomass crops, stream bank stabilization, living walls or snow fences, ornamental landscaping, and riparian buffers (Kuzovkina & Quigley, 2005; Stott, 1992). Evidence of sexual dimorphism for key traits suggests that sex of clones selected for particular uses can influence performance. This study demonstrates that *S. purpurea* expresses secondary sexual dimorphism for various traits where males in both a natural collection and in breeding populations responded more positively to multi-environmental and multi-year growing conditions. These findings also showed sex-specific differences in plasticity in response nitrogen amendments and disease pressure providing insights into resource allocation for primary versus reproductive growth, but whether there is sex-specific niche partitioning in shrub willow remains to be further evaluated. This study also determined that there is evidence of female biased sex ratio in *S. purpurea*. The results suggest that biased sex ratios in *S. purpurea* may be more strongly dependent on genetics, but recent advances in genomics will lead to genetic markers for early sex determination to help test this hypothesis.

## ACKNOWLEDGEMENTS

This work was supported by grants from the United States Department of Agriculture National Institute of Food and Agriculture through the Northeast Sun Grant Center (NE 11-48) and to the Northeast Woody/Warm-Season Biomass Consortium (NEWBio) (USDA-NIFA grant No. 2012-68005-19703), and also by a grant from the National Science Foundation Dimensions in Biodiversity Program (DEB-1542486). Junzhu Zou was supported by Chinese Scholarship Council (CSC). We thank all of those who provided technical assistance with field trial establishment, maintenance, and harvesting, especially: Curt Carter, Matt Christiansen, Brian DeGasperis, Lauren Carlson, Dawn Fishback, Steve Gordner, Michael Rosato, and Jeffrey Teague.

## AUTHOR CONTRIBUTIONS

F.E.G., C.H.C. and L.B.S. planned and designed the research. F.E.G. and C.H.C. wrote the first draft of the manuscript. F.E.G, C.H.C., L.B.S., J.Z., C.M.C., C.D.S., and L.M.E. performed the experiments, collected data, and analyzed the data. All authors reviewed and revised the final manuscript.

## SUPPORTING INFORMATION

**Appendix S1.** Clone ID, sex, and source information for 78 genotypes in the diversity panel.

**Appendix S2.** Experimental site characteristics for all trial locations.

**Appendix S3.** Materials and methods detailing genotyping, phenotypic and statistical analysis for *Salix purpurea*.

**Appendix S4.** Summary of phenotypic traits from the *Salix purpurea* diversity panel.

**Appendix S5.** Parameter estimates and significance values for multiple linear regression predictors of second year yield.

**Appendix S6.** Matrix of all pair-wise comparisons between traits by location within each year. The lower diagonal shows a scatter plot matrix with a LOESS smooth curve fitting, the main diagonal is a histogram showing the distribution of each trait, and the upper diagonal indicating the Pearson correlation coefficient (*r*) and *P*-value for each comparison. Locations: (A) Geneva, NY 2013 (B) Portland, NY 2013, (C) Morgantown, WV 2013, (D) Geneva, NY 2014, (E) Portland, NY 2014, and (F) Morgantown, WV 2014.

**Appendix S7.** Matrix of all pair-wise comparisons between traits measured in 2015 for the *Salix purpurea* F_1_ population.

**Appendix S8.** Matrix of all pair-wise comparisons between traits measured in 2015 for the *Salix purpurea* F_2_ population.

**Appendix S9** Correlation heatmap showing Pearson’s correlation coefficients (*r*) for all *Salix purpurea* accessions (n=78). Traits shown are divided by category as listed in Table 1. Colored boxes indicate significant correlations at *P*<0.05, where correlation coefficients of 1 are indicated by dark red and −1 shown as dark blue. All pair-wise comparisons between traits by year and location are shown in Appendix S6. Years: (A) 2013 and (B) 2014.

**Appendix S10.** Multiple linear regression model for estimating second year post-coppice biomass yield from annual measurements.

