## Supplemental Information S1-S10 for "Sexual dimorphism and sex ratio bias in the dioecious willow *Salix purpurea* L"

**Appendix S1** Clone ID, sex, and source information for 78 genotypes in the diversity panel.

| Clone ID | Epithet | Sex | Species | Latitude | Longitude | City/Town | State | Country | Notes |
|---|---|---|---|---|---|---|---|---|---|
| 94001 | | M | *S. purpurea* | N 43 13.000 | W 75 38 | Rome | NY | US | Natural accession |
| 94002 | | M | *S. purpurea* | N 43 13.001 | W 75 38 | Rome | NY | US | Natural accession |
| 94004 | | F | *S. purpurea* | N 43 13.003 | W 75 38 | Rome | NY | US | Natural accession |
| 94005 | | F | *S. purpurea* | N 43 13.004 | W 75 38 | Rome | NY | US | Natural accession |
| 94006 | | F | *S. purpurea* | N 43 13.005 | W 75 38 | Rome | NY | US | Natural accession |
| 94009 | | M | *S. purpurea* | N 42 52.002 | W 75 51 | Cazenovia | NY | US | Natural accession |
| 94011 | | F | *S. purpurea* | N 42 52.004 | W 75 51 | Cazenovia | NY | US | Natural accession |
| 94012 | | F | *S. purpurea* | N 42 52.005 | W 75 51 | Cazenovia | NY | US | Natural accession |
| 94013 | | F | *S. purpurea* | N 42 52.006 | W 75 51 | Cazenovia | NY | US | Natural accession |
| 94014 | | F | *S. purpurea* | N 42 52.007 | W 75 51 | Cazenovia | NY | US | Natural accession |
| 94015 | | F | *S. purpurea* | N 42 52.008 | W 75 51 | Cazenovia | NY | US | Natural accession |
| 95001 | | F | *S. purpurea* | N 42 51 | W 75 49 | New Woodstock | NY | US | Natural accession |
| 95002 | | M | *S. purpurea* | N 42 51.035 | W 75 46 | Nelson | NY | US | Natural accession |
| 95026 | | F | *S. purpurea* | N 41 46.3 | W 73 33 | Wassaic | NY | US | Natural accession |
| 95058 | | M | *S. purpurea* | N 42 18 | W 78 15 | Caneadea | NY | US | Natural accession |
| 00-01-001 | | M | *S. purpurea* | N 43 21.57 | W 75 32.6 | Lee | NY | US | Natural accession |
| 00-01-003 | | F | *S. purpurea* | N 43 21.585 | W 75 32.695 | Lee | NY | US | Natural accession |
| 00-01-004 | | F | *S. purpurea* | N 43 21.595 | W 75 32.758 | Lee | NY | US | Natural accession |
| 00-01-009 | | F | *S. purpurea* | N 43 23.986 | W 75 32.43 | Ava | NY | US | Natural accession |
| 00-01-011 | | F | *S. purpurea* | N 43 23.994 | W 75 32.43 | Ava | NY | US | Natural accession |
| 00-01-014 | | M | *S. purpurea* | N 43 10.06 | W 75 40.77 | Durhamville | NY | US | Natural accession |
| 00-01-034 | | F | *S. purpurea* | N 42 47.352 | W 76 11.434 | Tully | NY | US | Natural accession |
| 00-01-086 | | F | *S. purpurea* | N 42 53.639 | W 76 2.237 | Pompey | NY | US | Natural accession |
| 00-01-088 | | F | *S. purpurea* | N 42 49.493 | W 76 0.286 | Fabius | NY | US | Natural accession |
| 00-01-089 | | M | *S. purpurea* | N 42 50.325 | W 75 57.091 | Fabius | NY | US | Natural accession |
| 00-01-091 | | M | *S. purpurea* | N 42 44.682 | W 75 36.75 | Earlville | NY | US | Natural accession |
| 00-01-094 | | M | *S. purpurea* | N 42 43.997 | W 75 35.04 | Earlville | NY | US | Natural accession |
| 00-01-095 | | M | *S. purpurea* | N 42 33.185 | W 75 26.586 | New Berlin | NY | US | Natural accession |
| 00-01-098 | | F | *S. purpurea* | N 42 27.983 | W 75 25.426 | Mt. Upton | NY | US | Natural accession |

| | | | | | | | | | |
|---|---|---|---|---|---|---|---|---|---|
| 00-01-101 | | F | *S. purpurea* | N 42 54.168 | W 75 40.253 | Morrisville | NY | US | Natural accession |
| 00-01-102 | | M | *S. purpurea* | N 42 52.043 | W 75 40.82 | Eaton | NY | US | Natural accession |
| 00-01-103 | | F | *S. purpurea* | N 43 48.058 | W 75 38.653 | Lowville | NY | US | Natural accession |
| 00-01-104 | | F | *S. purpurea* | N 43 40.538 | W 75 26.473 | Turin | NY | US | Natural accession |
| 00-01-105 | | M | *S. purpurea* | N 43 34.208 | W 75 25.167 | Constableville | NY | US | Natural accession |
| 00-01-106 | | F | *S. purpurea* | N 43 21.942 | W 75 29.598 | Lee Center | NY | US | Natural accession |
| 01-01-001 | | M | *S. purpurea* | N 43 2.618 | W 76 3.021 | Fayetteville | NY | US | Natural accession |
| 01-01-028 | | F | *S. purpurea* | N 42 45.016 | W 76 7.893 | Tully | NY | US | Natural accession |
| 01-01-029 | | M | *S. purpurea* | N 42 54.388 | W 75 54.572 | Cazenovia | NY | US | Natural accession |
| 01-01-030 | | F | *S. purpurea* | N 42 54.365 | W 75 54.623 | Cazenovia | NY | US | Natural accession |
| 01-01-031 | | M | *S. purpurea* | N 42 54.386 | W 75 54.48 | Cazenovia | NY | US | Natural accession |
| 01-01-032 | | F | *S. purpurea* | N 42 53.42 | W 75 49.923 | Cazenovia | NY | US | Natural accession |
| 01-01-034 | | F | *S. purpurea* | N 42 53.183 | W 75 49.52 | Erieville | NY | US | Natural accession |
| 01-01-036 | | F | *S. purpurea* | N 42 51.235 | W 75 49.111 | New Woodstock | NY | US | Natural accession |
| 01-01-038 | | F | *S. purpurea* | N 42 48.12 | W 75 47.377 | Georgetown | NY | US | Natural accession |
| 01-01-042 | | M | *S. purpurea* | N 42 35.654 | W 75 35.141 | South Plymouth | NY | US | Natural accession |
| 01-01-078 | | M | *S. purpurea* | N 43 17.546 | W 75 16.081 | Holland Patent | NY | US | Natural accession |
| 01-01-079 | | M | *S. purpurea* | N 43 19.315 | W 75 19.673 | Westernville | NY | US | Natural accession |
| 01-01-082 | | F | *S. purpurea* | N 43 19.255 | W 75 19.924 | Westernville | NY | US | Natural accession |
| 01-01-084 | | F | *S. purpurea* | N 43 1.848 | W 76 8.528 | Syracuse | NY | US | Natural accession |
| 01-01-094 | | F | *S. purpurea* | N 42 33.767 | W 75 21.842 | New Berlin | NY | US | Natural accession |
| 01-07-251 | | F | *S. purpurea* | N 41 59.013 | W 73 21.315 | Canaan | CT | US | Natural accession |
| 02-201-005 | | M | *S. purpurea* | Location not recorded | | | | UKR | Natural accession |
| 03-01-005 | | F | *S. purpurea* | N 43 04.214 | W 76 15.649 | Solvay | NY | US | Natural accession |
| 03-01-007 | | F | *S. purpurea* | N 43 04.199 | W 76 15.649 | Solvay | NY | US | Natural accession |
| 03-01-013 | | F | *S. purpurea* | N 43 04.119 | W 76 15.689 | Solvay | NY | US | Natural accession |
| 03-01-017 | | F | *S. purpurea* | N 43 04.157 | W 76 15.791 | Solvay | NY | US | Natural accession |
| 03-01-019 | | F | *S. purpurea* | N 43 04.207 | W 76 15.819 | Solvay | NY | US | Natural accession |
| 03-01-020 | | F | *S. purpurea* | N 43 04.211 | W 76 15.740 | Syracuse | NY | US | Natural accession |
| 03-01-022 | | M | *S. purpurea* | N 43 04.309 | W 76 15.548 | Syracuse | NY | US | Natural accession |
| 03-01-023 | | F | *S. purpurea* | N 43 04.324 | W 76 15.425 | Syracuse | NY | US | Natural accession |
| 03-01-024 | | M | *S. purpurea* | N 43 04.337 | W 76 15.330 | Syracuse | NY | US | Natural accession |

| | | | | | | | | | |
|---|---|---|---|---|---|---|---|---|---|
| 03-01-025 | | M | *S. purpurea* | N 43 04.187 | W 76 14.256 | Syracuse | NY | US | Natural accession |
| 03-01-036 | | F | *S. purpurea* | N 43 48.299 | W 76 13.767 | Henderson | NY | US | Natural accession |
| 04-202-055 | ‘Denmark 601’ | M | *S. purpurea* | | | | | | Cultivar obtained from AgriGenesis |
| 04-202-058 | ‘Holland NZ 605’ | M | *S. purpurea* | | | | | | Cultivar obtained from AgriGenesis |
| 04-BN-046 | ‘Green Dicks’ | F | *S. purpurea* | | | | | | Cultivar obtained from Blue Stem Nursery |
| 05-01-001 | | M | *S. purpurea* | N 43 04.202 | W 76 15.645 | Syracuse | NY | US | Natural accession |
| 05-01-002 | | F | *S. purpurea* | N 43 04.212 | W 76 15.743 | Syracuse | NY | US | Natural accession |
| 05-01-003 | | F | *S. purpurea* | N 43 04.33 | W 76 15.187 | Syracuse | NY | US | Natural accession |
| 05-01-005 | | F | *S. purpurea* | N 43 04.240 | W 76 15.368 | Syracuse | NY | US | Natural accession |
| 05-OSU-063 | ‘Lambertiana’ | F | *S. purpurea* | | | | | US | Cultivar obtained from Ohio State University |
| 07-MBG-5095 | | F | *S. purpurea* | | | | | CAN | Natural accession obtained from Montreal Botanical Gardens |
| 07-MBG-5096 | | F | *S. purpurea* | | | | | CAN | Natural accession obtained from Montreal Botanical Gardens |
| 9882-34 | ‘Fish Creek’ | M | *S. purpurea* | | | | | US | Bred cultivar |
| 9882-41 | ‘Wolcott’ | F | *S. purpurea* | | | | | US | Bred cultivar |
| PMC9106302 | | F | *S. purpurea* | Location not recorded | | | | US | Natural accession obtained from USDA- NRCS |
| Pur12 | | M | *S. purpurea* | Location not recorded | | | | CAN | Natural accession obtained from University of Toronto |
| 05X-293-047 | | M | *S. purpurea* | | | | | US | Bred cultivar |

**Appendix S2** Experimental site characteristics for all trial locations.

| Site Characteristics[1] | Geneva, NY | | Portland, NY | Morgantown, WV |
|---|---|---|---|---|
| | $F_1$ & $F_2$ Trial | Association Trial | Association Trial | Association Trial |
| Latitude | 42°52’47”N | 42°52’11”N | 42°22'26"N | 39°39'31"N |
| Longitude | 77°00’55”W | 77°03’10”W | 79°29'11"W | 79°54'19"W |
| Elevation (m) | 184 | 234 | 228 | 365 |
| Soil Type | Odessa silt and lima loam | Lima Loam | Gravelly loam | Dormont and Guernsey silt loam |
| Nitrate (mg $kg^{-1}$) | - | 1.3 ± 0.7 | 10.2 ± 1.7 | 4.3 ± 2.1 |
| pH | - | 6.8 ± 0.0 | 5.1 ± 0.0 | 6.3 ± 0.2 |
| Organic (%) | - | 2.5 ± 0.0 | 3.8 ± 0.1 | 4.4 ± 0.3 |
| 2012 GDD[2] | 3041 | | 2990 | 3819 |
| 2013 GDD | 2731 | | 2654 | 3548 |
| 2014 GDD | 2678 | | 2499 | 3576 |
| 2015 GDD | 2730 | | 2835 | 2787 |
| 2012 Precipitation (May – August; cm) | 24.53 | | 30.40 | 29.31 |
| 2013 Precipitation (May – August; cm) | 46.38 | | 44.22 | 52.17 |
| 2014 Precipitation (May – August; cm) | 44.73 | | 42.09 | 49.56 |
| 2015 Precipitation (May – August; cm) | 40.03 | | 13.21 | 33.17 |

[1]Weather data for Geneva and Portland, NY were obtained from Cornell University’s Network for Environment and Weather Applications database. Weather data for Morgantown, WV was collected from the National Oceanic and Atmospheric Administration website. Means ± standard error are shown for nitrate, pH , and organic matter

[2]GDD; growing degree days

**Appendix S3 Materials and Methods**

Phenotyping

*Morphology and Biomass*

During the dormant period after each growing season, stem diameter (cm) and stem number was measured. The diameters of stems greater than or equal to 5 mm was measured at 30 cm from the base of the plant using Masser Racal 500 digital calipers (Masser, Rovaniemi, Finland). Stem area ($cm^2$) was also calculated using the stem diameter values. Maximum stem height (m) of every plot was recorded using a measuring rod (Crain Enterprises, Inc., Mound City, IL), and the mean height was calculated for each plot. In July of each year, internode length (cm) was measured within the middle third on the tallest stem of each plot and the length of five internodes were recorded. Accounting for different phyllotactic patterns, alternate leaves were counted using five alternate buds or leaves from the first designated bud/leaf, whereas opposite leaves or buds were counted as one node. At the end of the second growing season, crown diameter (cm) was measured using modified Haglöf Mantax blue forestry calipers (Haglöf Sweden AB, Långsele, Sweden). Stool diameters were measured at 15 cm above the soil, which is the average height of a shrub willow harvester. Crown form (branching angle) was calculated by using one-half of the crown diameter measurement and the height at which it was measured (15 cm) to find the angle of the stem branching relative to the soil. Leaf perimeter (cm), maximum leaf length (cm), leaf width (cm) and leaf area ($cm^2$) were measured on mature leaves at mid-canopy level on the tallest stem of each plant per plot using a laser leaf area meter (CI-203 model, CID Bio-Science, Inc., USA). The same measurement leaves were collected and completely dried at 65°C and weighed. The dry weight and measured leaf area were used to calculate specific leaf area (SLA) ($cm^2\ g^{-1}$ dry wt).

For the diversity panel, yield was measured after the second year of post-coppice by harvesting and weighing each four plant plot using the Ny Vraa JF192 harvester (Ny Vraa Bioenergy, Tylstrup, Denmark). Chips were collected in a plastic bin mounted on Avery Weigh-Tronix weigh cells (Fairmont, MN), and the wet weight of the chip biomass of each plot was recorded. A sub-sample of fresh chip biomass (1 kg) was collected for each plot, weighed after harvest, oven-dried at 65ºC to a constant weight, and dry weight recorded to determine moisture content at harvest. The chip dry weight was then used to estimate plot dry weights from the measured fresh weights. For all plots, dry biomass yield was calculated and expressed in dry Mg $ha^{-1}$ based on plot area.

*Phenology*

Floral and vegetative bud break were observed and scored using a 0-5 rating scale only in the second year of growth due to the absence of floral buds in the first year. The established scale used for phenology ratings was modified from Saska *et al.* (2010). Both floral and vegetative phenology was surveyed once a week for five weeks and was recorded as the day of the year for a given rating that was observed. All observations occurred until all stage 5 scores were recorded for every genotype. For all three populations, the sex of each genotype was visually scored and recorded as either male (M), female (F), or hermaphrodite (H).

*Physiology*

Stomatal conductance ($g_s$) (mmol $m^{-2}s^{-1}$) was measured on the abaxial side of the leaf with a leaf porometer (SC-1 Leaf Porometer, Decagon, Pullman, WA) on the uppermost fully expanded leaf of the tallest stem of the plant. A non-destructive proxy for monitoring nitrogen status in the plant was measured by relative chlorophyll content (SPAD) with a portable chlorophyll meter (SPAD-502, Minolta Osaka Co., Ltd., Japan) where three measurements were taken along the length of the tallest stem from the upper, middle, and lower canopy levels and averaged for each plot.

The canopy color (RGB-15) of $F_1$ and $F_2$ families was determined using aerial images collected with a gimbal-mounted 14 Megapixel F/2.8 140° FOV camera (w/ lens stabilization) of a Phantom 2 Vision+ (DJI, Nanshan District, Shenzhen, China) quadcopter. To account for any variation, three replicate images were taken for each interval at a fixed altitude (120 ft) along the length of the field trial (1,196 ft) in late-July 2015. An overlap of each interval was required to properly interleave them into a single image. Images were lens corrected using the DJI Vision plugin, ordered, and interleaved using the in Adobe Photoshop C6 (Adobe Systems Incorporated, San Jose, CA). The resulting interleaved full-field images were converted into separate RGB channels and analysed using a colorimetric scale based on green pixel density in the open-source program ImageJ v1.47 (Rasband, 1997-2016; Schneider *et al.*, 2012). Excluding aisles and border plants, a coordinate grid of the field was developed in order to obtain average pixel density for each plot.

*Wood Properties*

For analysis of physical and chemical wood properties in the diversity panel, stem segment samples were collected after each growing season in the dormant period. Stem segments were collected based on sampling methods previously described (Liu *et al.*, 2015) and were stored frozen at - 3°C until they were processed. The specific gravity of each sample was measured by volumetric displacement (om-06, 2006). In 2014, a modified method of measuring specific gravity was used where the volume of water displaced was weighed for added precision. Following specific gravity determination, stem segments were oven-dried at 65°C to a constant weight and then rough milled to a 5 mm particle size with a Retch SM300 cutting mill (Retch, Haa, Germany) and were further comminuted to a 0.5 mm particle size by fine milling with the IKA MF 10.1 knife mill (IKA, Wilmington, NC) for compositional analysis. Approximately 20 mg of each milled, unextracted stem sample was analyzed with a Thermogravimetric Analyzer Q500 instrument and Universal Analysis 2000 ver. 4.5A software (TA Instruments, New Castle, DE), as previously described (Serapiglia *et al.*, 2009). Hemicellulose, cellulose, lignin, and ash content were then determined as a percentage of total dry biomass for each sample as previously described (Serapiglia *et al.*, 2014).

*Disease Severity Assessment*

In September 2015, natural infection of *Melampsora* spp. was visually scored in each population based on percentage of leaf area covered by rust pustules. Percent disease severity was scored (0-100%) for each plot. For the diversity panel, defoliation occurred when a leaf was 50% covered with rust pustules, thus the highest degree recorded in this population was 50%.

**Appendix S4 -** Summary of phenotypic traits from the *S. purpurea* diversity panel

| Trait[1] | Units | Mean (±SE) | Min. – Max. |
|---|---|---|---|
| **Biomass** | | | |
| Height-13 | m | 1.92±0.01 | 0.11-3.26 |
| Height-14 | m | 3.14±0.02 | 0.4-4.88 |
| Stem Number-13 | # | 17.95±0.27 | 2-54 |
| Stem Number-14 | # | 21.92±0.31 | 0-73 |
| Mean Stem Diameter-13 | cm | 7.33±0.04 | 2-12.73 |
| Mean Stem Diameter-14 | cm | 10.05±0.27 | 3.6-17.71 |
| Total Stem Area-13 | $cm^2$ | 9.3±0.19 | 0.13-46.6 |
| Total Stem Area-14 | $cm^2$ | 21.04±0.37 | 0.13-84.38 |
| Internod Length-13 | cm | 13.51±0.14 | 3.5-57 |
| Internod Length-14 | cm | 13.36±0.18 | 4.55-32 |
| Yield-14 | dry Mg $ha^{-1}$ | 2.74±0.05 | 0.11-16.51 |
| **Foliar** | | | |
| Leaf Length-13 | cm | 6.95±0.05 | 1.25-15.87 |
| Leaf Length-14 | cm | 6.72±0.05 | 1.36-16.17 |
| Leaf Width-13 | cm | 1.75±0.04 | 0.46-19.64 |
| Leaf Width-14 | cm | 1.9±0.05 | 0.69-25.73 |
| Leaf Area-13 | $cm^2$ | 9.06±0.15 | 0.58-50.55 |
| Leaf Area-14 | $cm^2$ | 9.01±0.11 | 1.04-37.47 |
| Leaf Perimeter-13 | cm | 24.01±0.8 | 2.82-197.6 |
| Leaf Perimeter-14 | cm | 17.83±0.23 | 2.76-68.89 |
| Leaf Weight-13 | g | 0.08±0.001 | 0.01-0.73 |
| Leaf Weight-14 | g | 0.1±0.001 | 0.02-0.4 |
| Specific Leaf Area-13 | $cm^2\ g^{-1}$ | 129.41±3.07 | 6.66-1713.56 |
| Specific Leaf Area-14 | $cm^2\ g^{-1}$ | 91.64±0.55 | 22.63-314.73 |
| **Architecture** | | | |
| Crown Diameter-14 | cm | 29.96±0.42 | 1-183.7 |
| Crown Form-14 | ° | 48.71±0.38 | 9.4-88.1 |
| **Composition** | | | |
| Hemicellulose-13 | % | 17.59±0.03 | 14.41-21.06 |
| Hemicellulose-14 | % | 17.73±0.03 | 15.87-21.77 |
| Cellulose-13 | % | 37.55±0.11 | 25.91-47.3 |
| Cellulose-14 | % | 41.56±0.06 | 33.27-45.89 |
| Lignin-13 | % | 28.9±0.07 | 24.37-36.98 |
| Lignin-14 | % | 27.32±0.04 | 24.17-32.38 |
| Ash-13 | % | 2.17±0.02 | 0.78-4.31 |
| Ash-14 | % | 1.57±0.01 | 0.82-3.57 |
| Specific Gravity-13 | g $cm^{-3}$ | 0.45±0.002 | 0.23-0.79 |
| Specific Gravity-14 | g $cm^{-3}$ | 0.49±0.001 | 0.32-0.73 |
| **Physiology** | | | |
| AugSPAD-13 | SPAD units | 45.86±0.25 | 17.1-75 |

| | | | |
|---|---|---|---|
| SeptSPAD-13 | SPAD units | 45.38±0.19 | 28.8-69.9 |
| AugSPAD-14 | SPAD units | 41.6±0.29 | 17.2-75.1 |
| SeptSPAD-14 | SPAD units | 42.92±0.22 | 20.33-91 |
| Stomatal Conductance-13 | mmol $m^{-2} s^{-1}$ | 594.73±5.44 | 45.5-1164.2 |
| Stomatal Conductance-14 | mmol $m^{-2} s^{-1}$ | 477.91±4.47 | 80.8-914.1 |
| **Phenology** | | | |
| Vegetative Phenology-14 | day of year | 109.92±0.12 | 102-118 |
| Floral Phenology-14 | day of year | 92.17±0.49 | 57-115 |

[1]Phenotypic traits measured in years 2013 (-13) and 2014 (-14). See Appendix S3 for trait definitions.

**Appendix S5** Parameter estimates and significance values for multiple linear regression predictors of second year yield.

| **Variable**[1] | **Estimate** | ***t*** | ***P*-value** | **95% Confidence Limits** | |
|---|---|---|---|---|---|
| Intercept ($\beta_0$) | -2.56 | -12.24 | <.0001 | -2.97 | -2.15 |
| SA-13 ($\beta_1$) | 0.06 | 8.96 | <.0001 | 0.05 | 0.08 |
| SA-14 ($\beta_2$) | 0.92 | 20.95 | <.0001 | 0.83 | 1.00 |
| HT-14 ($\beta_3$) | 0.05 | 13.59 | <.0001 | 0.04 | 0.06 |
| AugSPAD-14 ($\beta_4$) | 0.01 | 3.48 | 0.0005 | 0.01 | 0.02 |

[1]Significant predictor variables measured in 2013 (-13) and 2014 (-14).

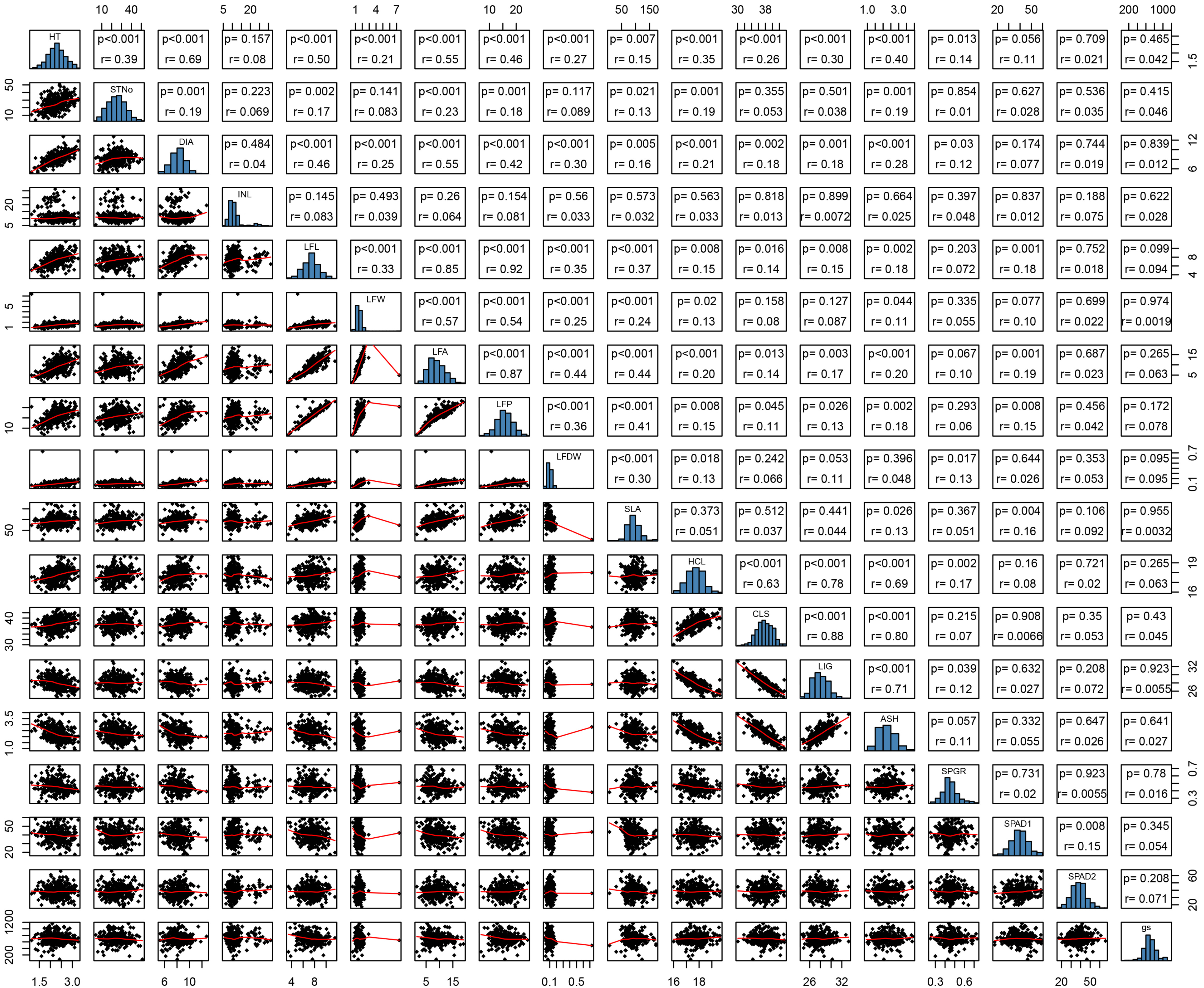


**Appendix S6**. Matrix of all pair-wise comparisons between traits by location within each year. The lower diagonal shows a scatter plot matrix with a LOESS smooth curve fitting, the main diagonal is a histogram showing the distribution of each trait, and the upper diagonal indicating the Pearson correlation coefficient (r) and P-value for each comparison. Locations: (A) Geneva, NY 2013

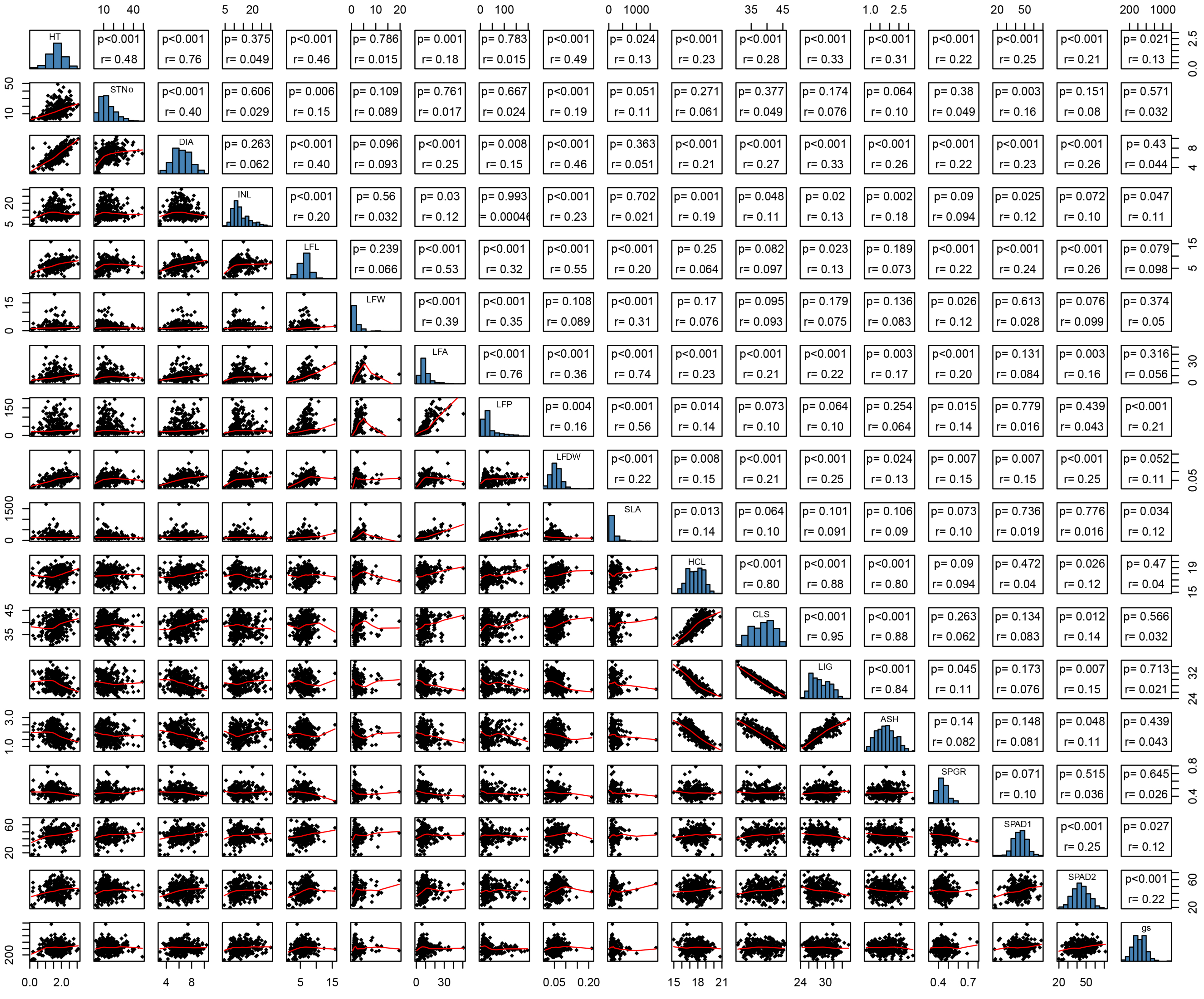

**Appendix S6.** Matrix of all pair-wise comparisons between traits by location within each year. The lower diagonal shows a scatter plot matrix with a LOESS smooth curve fitting, the main diagonal is a histogram showing the distribution of each trait, and the upper diagonal indicating the Pearson correlation coefficient (r) and P-value for each comparison. Locations: (B) Portland, NY 2013

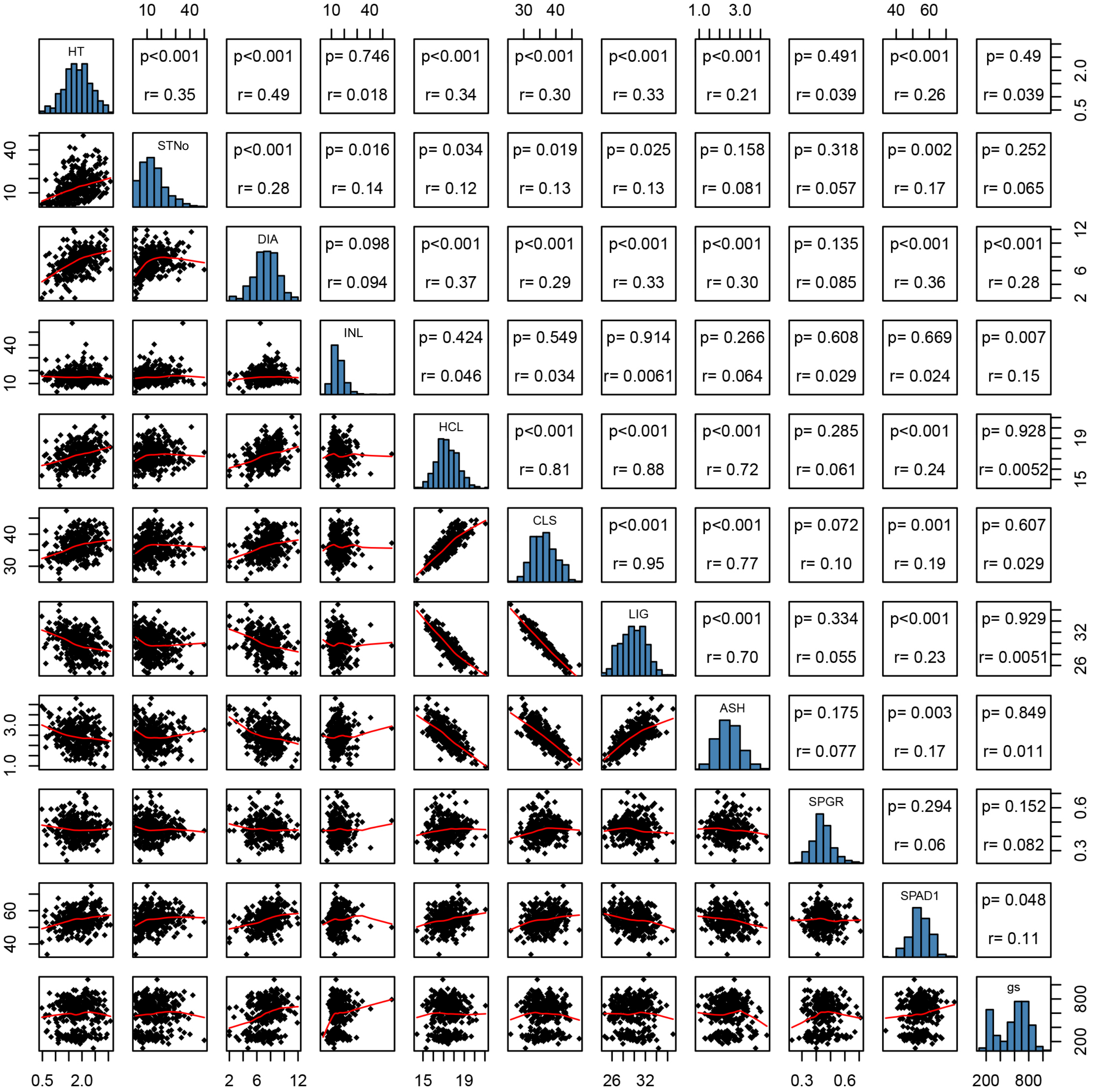


**Appendix S6**. Matrix of all pair-wise comparisons between traits by location within each year. The lower diagonal shows a scatter plot matrix with a LOESS smooth curve fitting, the main diagonal is a histogram showing the distribution of each trait, and the upper diagonal indicating the Pearson correlation coefficient (r) and P-value for each comparison. Locations: (C) Morgantown, WV 2013

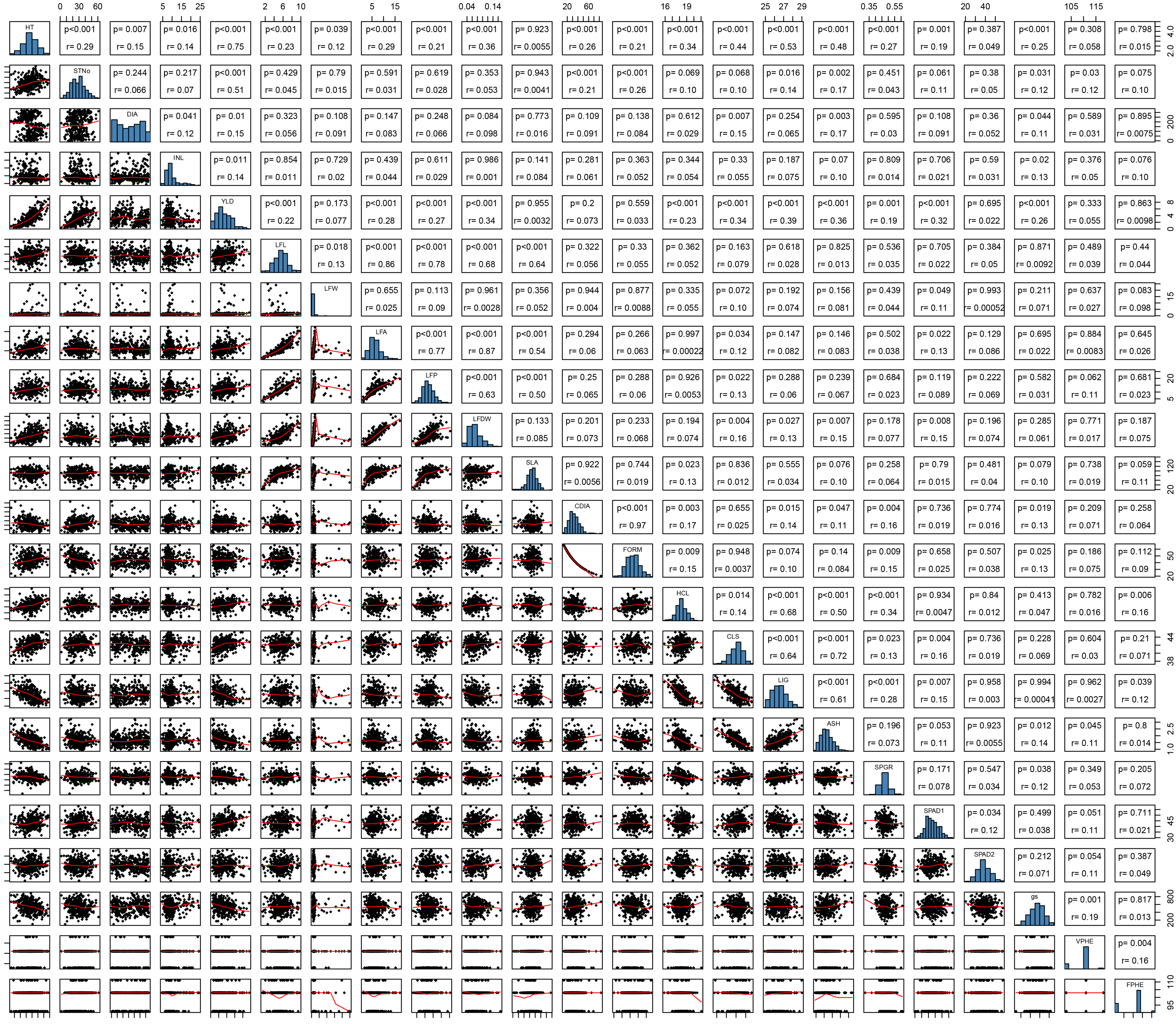


**Appendix S6.** Matrix of all pair-wise comparisons between traits by location within each year. The lower diagonal shows a scatter plot matrix with a LOESS smooth curve fitting, the main diagonal is a histogram showing the distribution of each trait, and the upper diagonal indicating the Pearson correlation coefficient (r) and P-value for each comparison. Locations: (D) Geneva, NY 2014

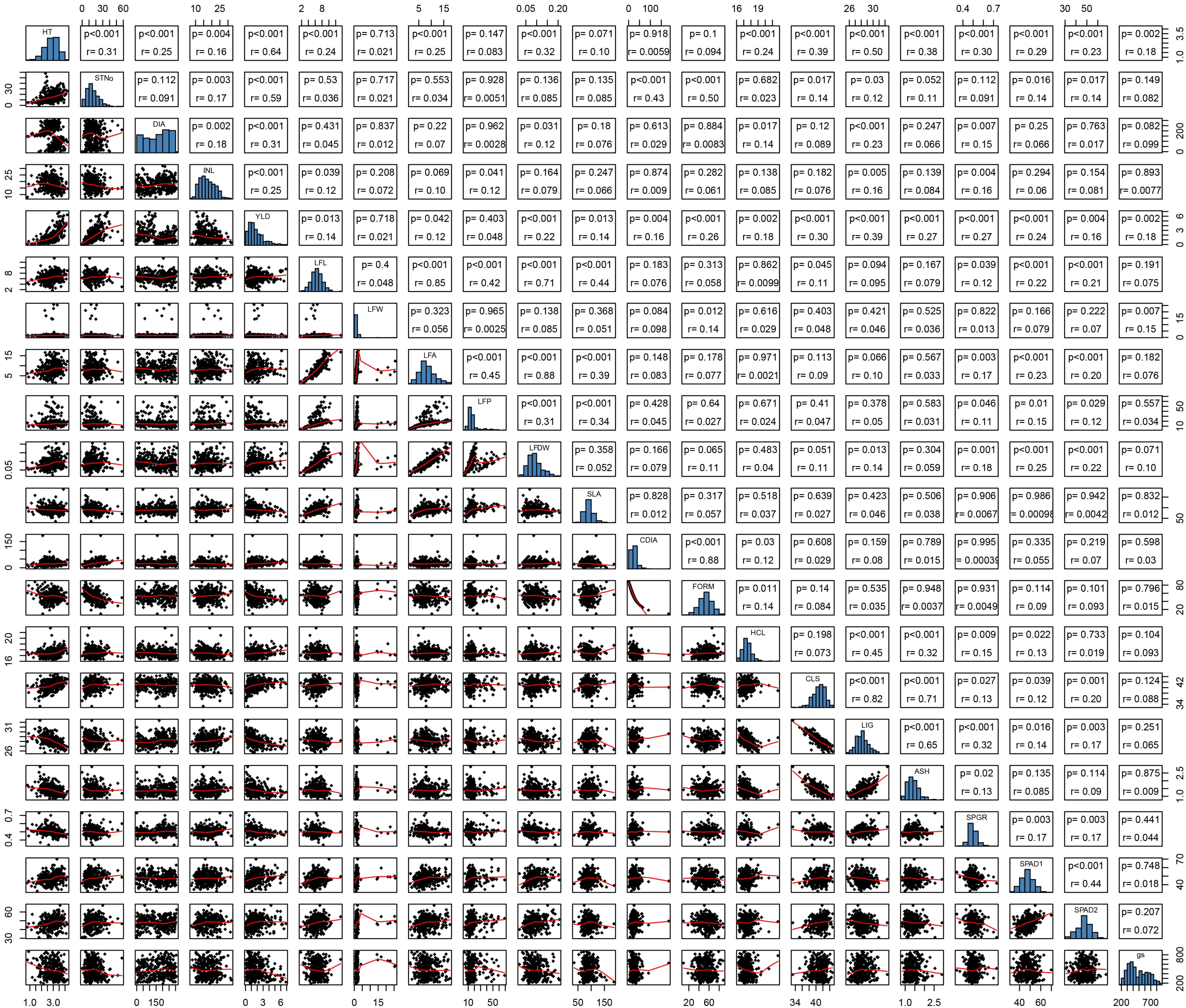


**Appendix S6**. Matrix of all pair-wise comparisons between traits by location within each year. The lower diagonal shows a scatter plot matrix with a LOESS smooth curve fitting, the main diagonal is a histogram showing the distribution of each trait, and the upper diagonal indicating the Pearson correlation coefficient (r) and P-value for each comparison. Locations: (E) Portland, NY 2014

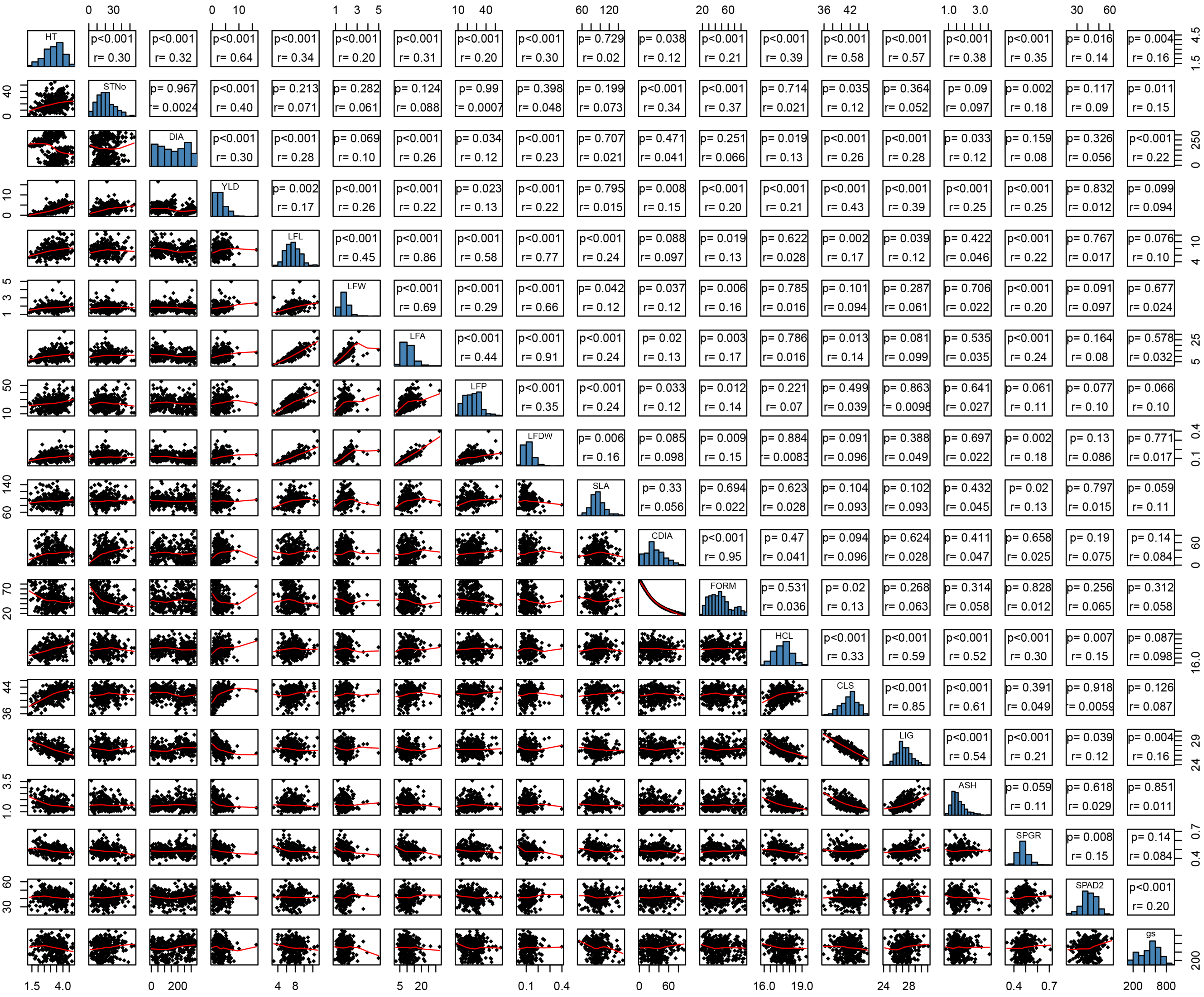


**Appendix S6.** Matrix of all pair-wise comparisons between traits by location within each year. The lower diagonal shows a scatter plot matrix with a LOESS smooth curve fitting, the main diagonal is a histogram showing the distribution of each trait, and the upper diagonal indicating the Pearson correlation coefficient (r) and P-value for each comparison. Locations: (F) Morgantown, WV 2014

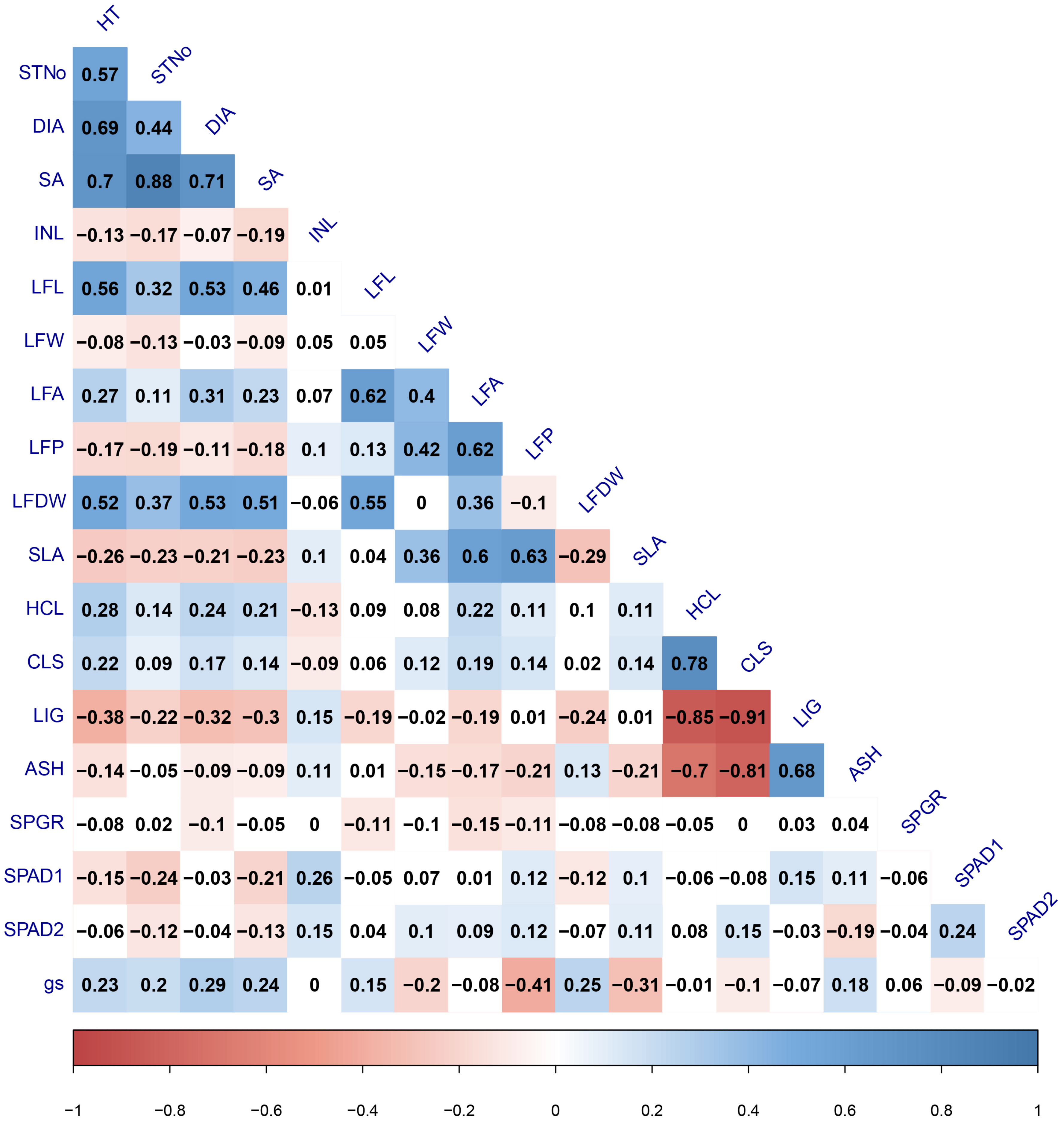


**Appendix S7**. Matrix of all pair-wise comparisons between traits measured in 2015 for the *Salix purpurea* $F_1$ population.

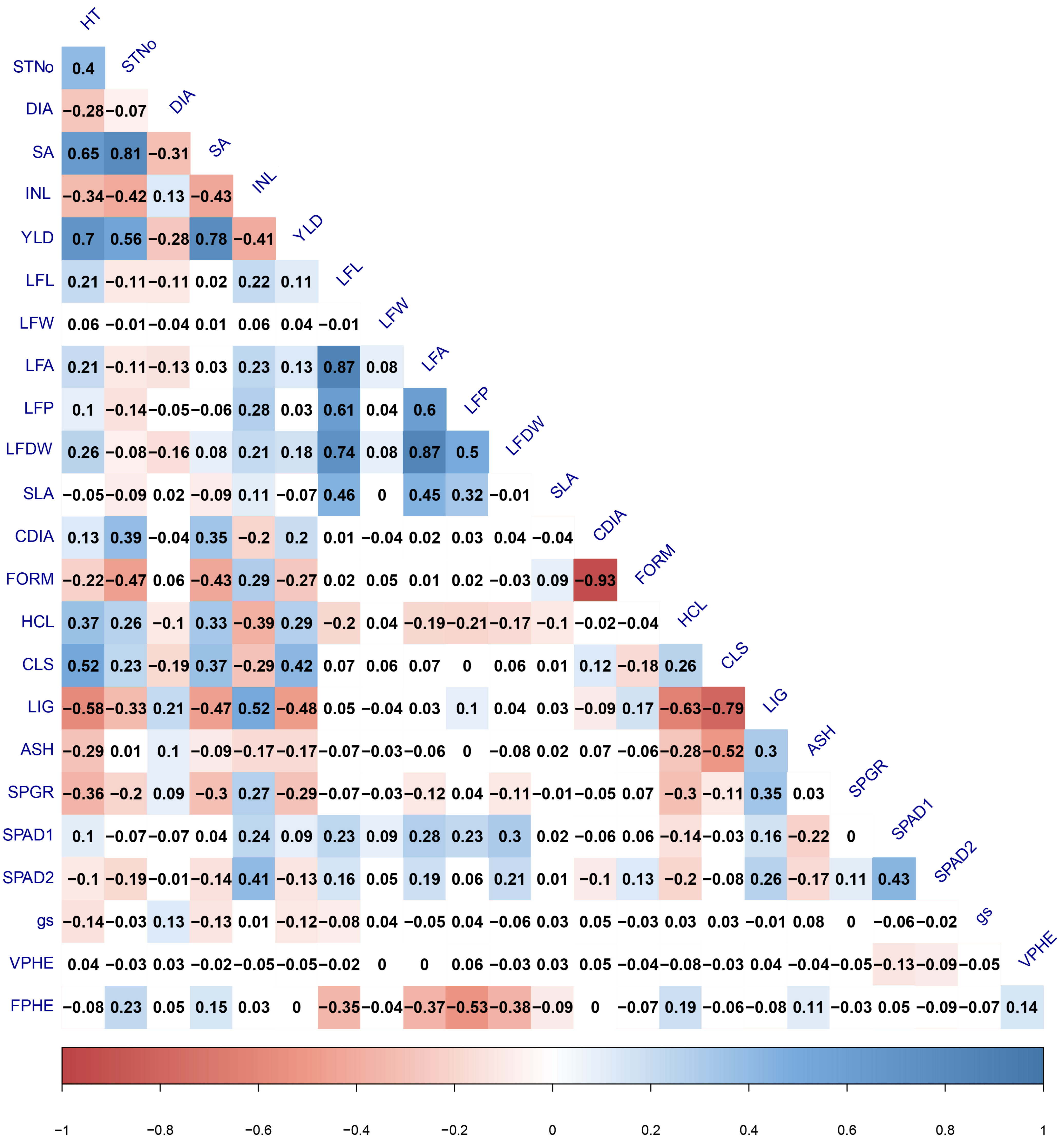


**Appendix S8**. Matrix of all pair-wise comparisons between traits measured in 2015 for the *Salix purpurea* $F_2$ population.

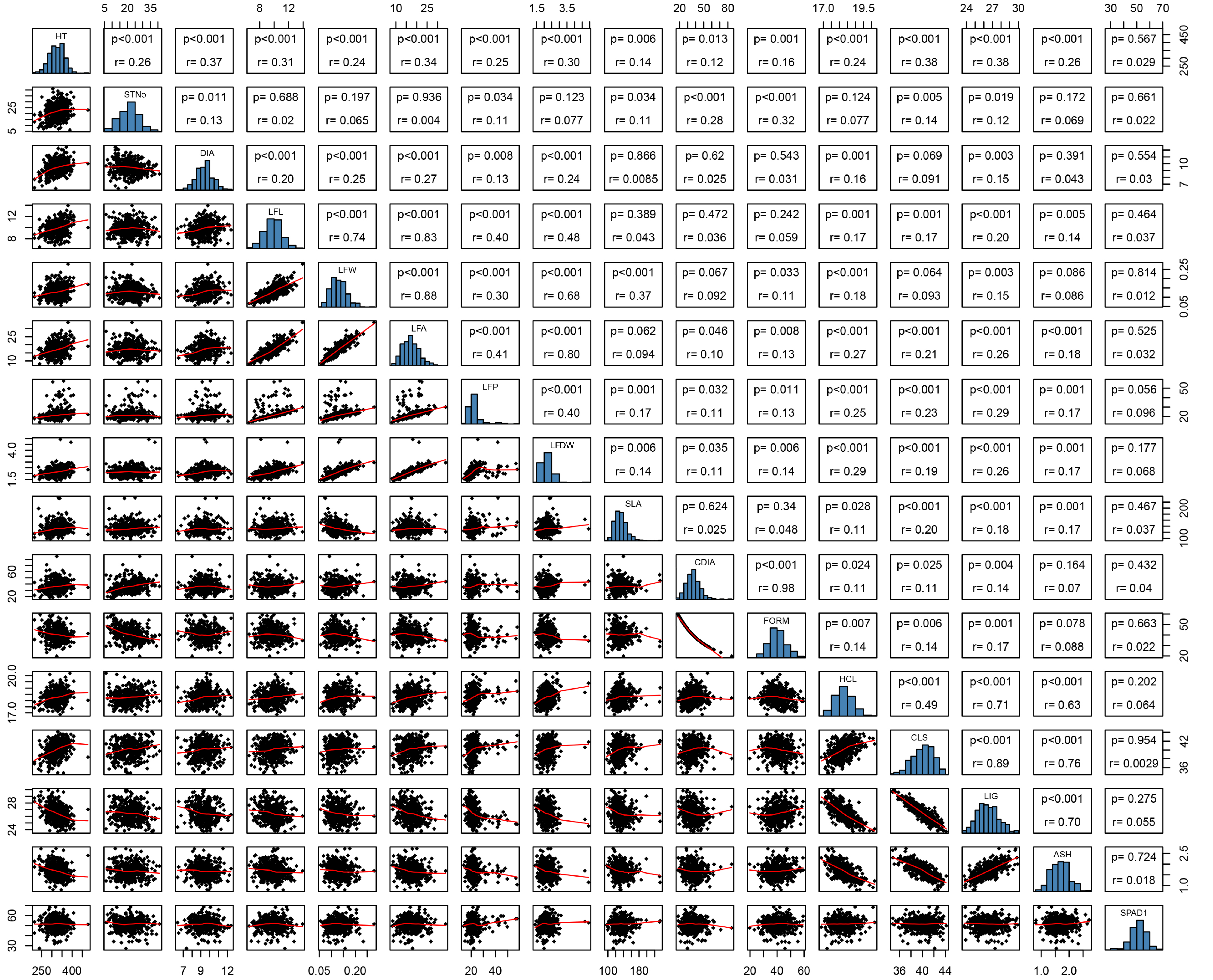


**Appendix S9** Correlation heatmap showing Pearson's correlation coefficients (r) for all *Salix purpurea* accessions (n=78). Traits shown are divided by category as listed in Table 1. Colored boxes indicate significant correlations at P<0.05, where correlation coefficients of 1 are indicated by dark red and -1 shown as dark blue. All pair-wise comparisons between traits by year and location are shown in Appendix S6. Years: (A) 2013

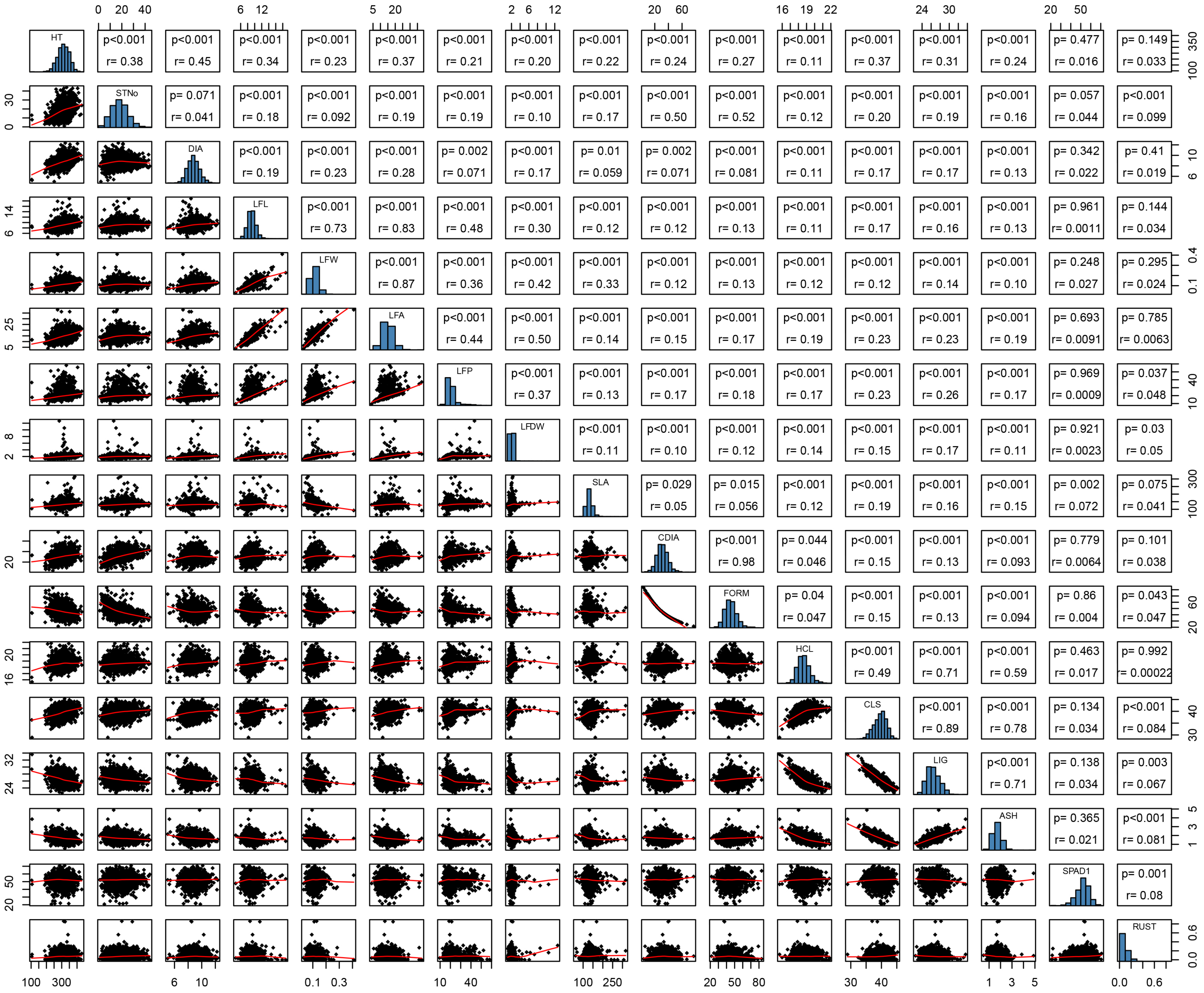


**Appendix S9** Correlation heatmap showing Pearson's correlation coefficients (r) for all Salix purpurea accessions (n=78). Traits shown are divided by category as listed in Table 1. Colored boxes indicate significant correlations at P<0.05, where correlation coefficients of 1 are indicated by dark red and -1 shown as dark blue. All pair-wise comparisons between traits by year and location are shown in Appendix S6. Years: (B) 2014

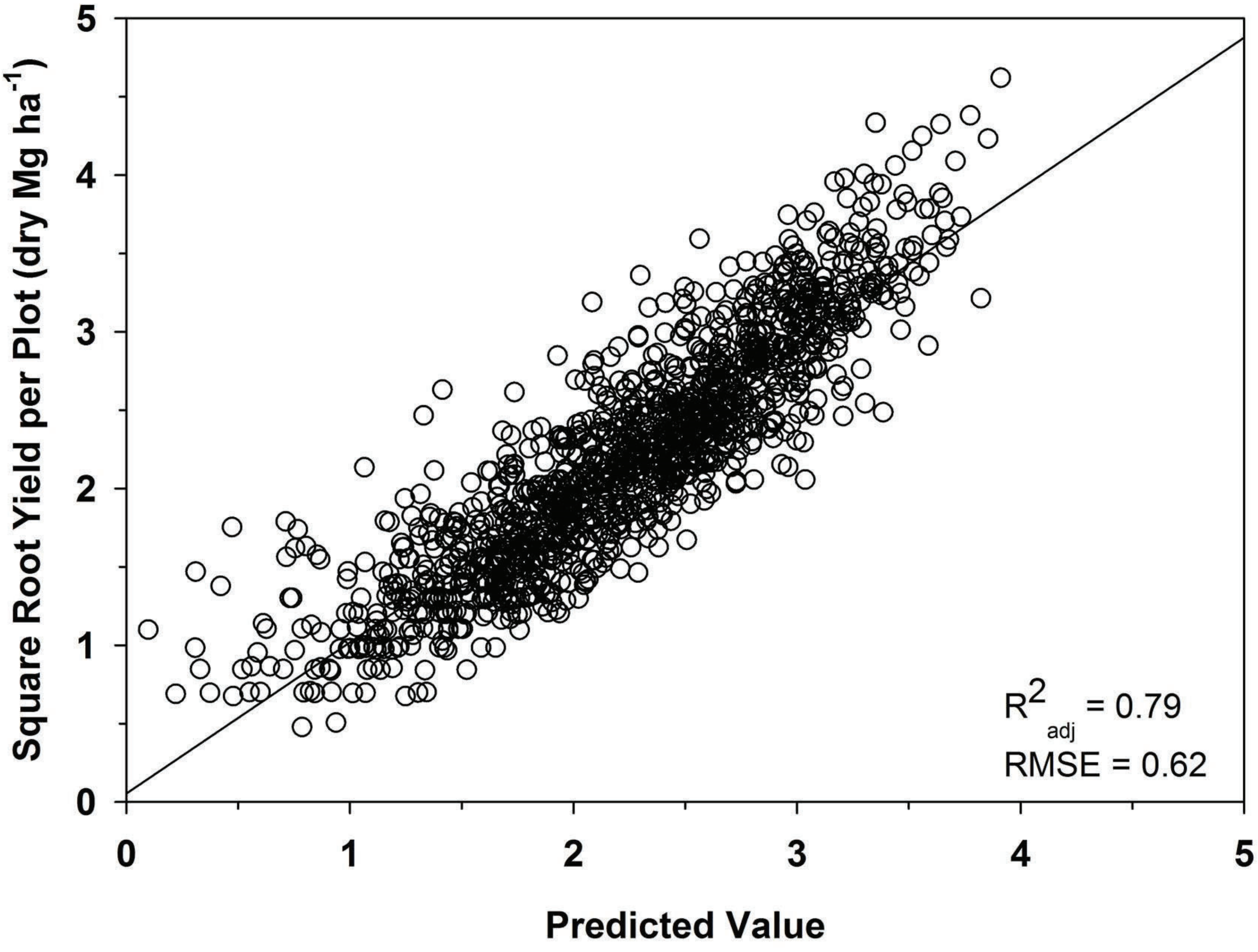


**Appendix S10**. Multiple linear regression model for estimating second year post-coppice biomass yield from annual measurements.
